# Preclinical modeling of surgery and steroid therapy for glioblastoma reveals changes in immunophenotype that are associated with tumor growth and outcome

**DOI:** 10.1101/2020.08.12.248443

**Authors:** Balint Otvos, Tyler J. Alban, Mathew M. Grabowski, Defne Bayik, Erin E. Mulkearns-Hubert, Tomas Radivoyevitch, Anja Rabljenovic, Sarah Johnson, Charlie Androjna, Alireza M. Mohammadi, Gene Barnett, Manmeet S. Ahluwalia, Michael A. Vogelbaum, Peter E. Fecci, Justin D. Lathia

## Abstract

Recent advances in cancer immunotherapy have created a greater appreciation of potential anti-tumoral impacts by the immune system; however, individual patient responses have been variable. While immunotherapy is often given after standard-of-care treatment, the effects of initial interventions on the ability of the immune system to mount a response are not well understood and this may contribute to the variable response. For glioblastoma (GBM), initial disease management includes surgical resection, perioperative high-dose steroid therapy, chemotherapy, and radiation treatment. While new discoveries regarding the impact of chemotherapy and radiation on immune response have been made and translated to clinical trial design, the impact of surgical resection and steroids on the anti-tumor immune response has yet to be determined. Further, it is now accepted that steroid usage needs to be closely evaluated in the context of GBM and immunotherapy trials. To better model the clinical scenario in GBM, we developed a mouse model that integrates tumor resection and steroid treatment to understand how these therapies affect local and systemic immune responses. Using this model, we observed a systemic reduction in lymphocytes associated with surgical resection and identified a correlation between increased tumor volume and decreased circulating lymphocytes, a relationship that was obviated by dexamethasone treatment. Furthermore, we investigated the possibility of there being similar relationships in a cohort of patients with GBM and found that prior to steroid treatment, circulating lymphocytes inversely correlated with tumor volume. Lastly, correlating GBM patient data and outcomes demonstrated that peripherally circulating lymphocyte content varies with progression-free and overall survival, independent of tumor volume, steroid use, or tumor molecular profiles. These results highlight the systemic immunosuppressive effects that initial therapies can have on patients. Such effects should be considered when designing current and future immunotherapy clinical trials and underscore the importance of circulating lymphocytes as a possible correlate of GBM disease progression.

## Introduction

Glioblastoma (GBM), a WHO grade IV glioma, is treated with standard-of-care therapies including maximal safe surgical resection, steroids, and concomitant radiation and chemotherapy with temozolomide. Despite the short-term efficacy of these approaches, the 5-year survival rate remains only 5%^1^. Following the recent success of immunotherapy in cancers such as high-grade melanoma and lung cancer, there is currently an extensive clinical trial effort to produce similar anti-tumor immune responses in patients with GBM^2^. Thus far, these trials have largely failed to impact survival in GBM patients, but they have successfully drawn attention to the potent local and systemic immune suppression elicited by GBM^2–6^. In seeking to develop better immune therapies for GBM, many have logically sought to understand how standard-of-care chemotherapy^7–10^ and radiation^11–13^ impact the immune system and the tumor immune microenvironment. Although neoadjuvant immunotherapy prior to surgical resection may lead to more effective treatment responses^14^, the impact of standard surgical resection and steroid treatment otherwise remains largely unexplored. This lack of understanding is in part because almost every patient who undergoes surgical resection also receives steroid treatment to reduce edema, confounding the study of singular effects in patients. In addition, patients typically undergo chemotherapy and radiation shortly after surgery, adding to the difficulty in understanding how surgery and steroids impact the anti-tumor immune response.

To develop a better understanding of the immunological changes that occur due to standard-of-care treatment, we developed a mouse model of surgical resection that also employs a clinically relevant steroid dose. We utilized two syngeneic murine GBM models (GL261 and CT-2A^15^) and found that surgical resection alone elicited a subsequent reduction in peripheral T cells, as did treatment with steroids. Additionally, we identified a negative correlation between the number of T cells in the circulation and tumor volume. We confirmed similar correlations in a GBM patient cohort, where patients who did not receive steroid treatment exhibited a negative correlation between peripheral blood lymphocyte counts and tumor volume. Initial lymphocyte count was a significant predictor of progression-free- and overall survival in multivariate models.

## Results

### Establishment of a clinically relevant mouse model of GBM surgical resection

Given the inherent difficulties of studying the singular effects of surgical resection or steroid treatment in patients with GBM, we sought to develop a murine model of GBM surgical resection with a translationally relevant steroid dose. The steroid-treated mice received 4 μg dexamethasone daily, which corresponds by weight to a dosage of 16 mg of dexamethasone per day for an 80 kg patient. In this initial proof-of-concept investigation, we first intracranially injected GL261 cells at day 0 and waited 7 days before surgically resecting the tumor or performing a mock resection procedure, in which a corticectomy was performed after opening the dura and separating white matter tracks overlying the tumor without removing the tumor. The mock resection cohort was developed to normalize for systemic murine responses to injury or violation of the dura/cranial cortical surface (**Fig. 1A**)^16^. Importantly, time under anesthesia, blood loss, and surgical procedure time did not differ significantly between the two cohorts: the mean time of the surgical procedure was ~20 minutes in both cohorts. MRI images were obtained for each mouse prior to surgical intervention, after resection, and 7 days post-procedure, when recurrent tumor spreading throughout the brain was typically observed, usually either along the surface of the resection cavity or along white matter tracts (**Fig. 1B**). Volumetric analysis of post-resection animals, in both the PBS and dexamethasone 4 μg daily cohorts, demonstrated successful tumor debulking and resultant decreases in recurrent tumor volumes at 7 days post-surgery (**Fig. 1C, D, Supplemental Figure 1A**). Utilizing this model, surgical resections compared to mock surgical corticectomies demonstrated a median survival benefit of 4 days (**Fig. 1E**). As expected, administration of therapeutic levels of dexamethasone (4 μg daily per 20 g animal) did not alter survival in either GL261 or CT-2A tumor-bearing mice with or without surgical resection of the tumors (**Fig. 1F, Supplemental Fig. 1B, C**). Importantly, MRI analysis of dexamethasone-treated mice showed the expected reduction in tumor-induced vasogenic edema (**Fig. 1G, H**). These data demonstrate a clinically appropriate model of surgical resection and steroid treatment in mice that can be utilized to investigate immunological changes due to initial upfront GBM therapies.

**Figure 1.**
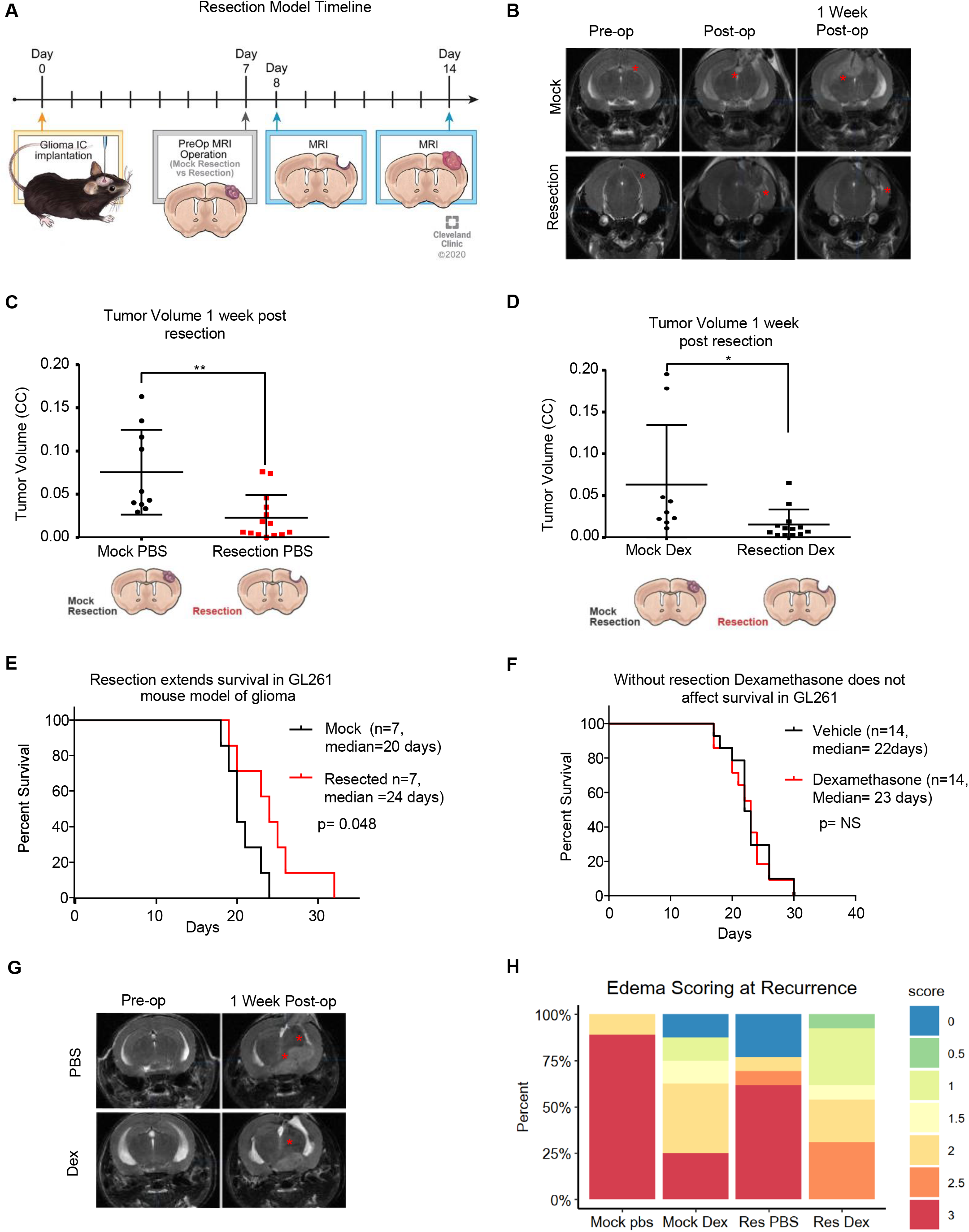
Murine model of resection including dexamethasone treatment extends survival and reduces edema. To replicate standard-of-care and dexamethasone treatment, a mouse model of resection was initiated as outlined in the diagram, with intracranial implantation of tumor, followed by MRI prior to resection and after resection, and then endpoint MRI along with flow cytometry or survival depending on the experiment (**A**). Representative MRI images of tumors from the mock resection and resection cohorts pre-, post- and 1 week post-resection (**B**). Tumor volume was assessed using the BrainLab software suite and graphed as tumor volume (**C, D**). A survival study comparing n=7 mice with mock resection and n=7 mice with resection was performed showing a median survival of 20 days for mock resection vs 24 days for resection, with log-rank p-value shown (**E**). Vehicle-vs dexamethasone (one week at 4 μg daily)-treated GL261-bearing mice with no surgical resection were also compared and showed no survival difference due to dexamethasone (**F**). Representative MRI images of dexamethasone-treated mice and PBS-treated mice pre-op and 1 week post-op (**G**). Edema scoring is graphed as a percentage with n=7 mice per group (**H**). Student’s two-tailed t-test was performed for comparisons in panels A, D, E; *p<0.05, **p<0.01, ***p<0.001. Survival curve analysis was performed in GraphPad Prism using log-rank tests (also known as Mantel-Cox tests) for p values.

### Surgical resection causes systemic immune suppression

After development of the surgical resection model, we sought to utilize this model to determine whether there are differences in local and systemic immune responses related to initial treatment. For studies relating to the immunobiology of treatment, we chose a time point of 7 days post-resection to allow for the effects of dexamethasone treatment to develop and for temporal resolution of the presumed initial injury and healing response to surgery. This also permitted analysis of the recurrent tumor at that time. Based on previous studies of CNS injury response^16^, this time point should show the lasting immunologic effects separate from the acute injury response. Thus, mice in an initial cohort were implanted with tumors, that were allowed to grow before being resected as described above, and 7 days post-resection, an MRI was performed. Flow cytometry for lymphoid (T cells, CD4+ T cells, CD8+ T cells, natural killer (NK) cells, T regulatory cells) and myeloid (myeloid-derived suppressor cells (MDSCs), macrophages, dendritic cells, and monocytes) cells was performed on samples of blood, spleen, gross recurrent tumor, non-tumor cortex within the recurrent tumor-bearing hemisphere, and bone marrow. There were no major differences noted between the percentage of CD45+ live cells present among any of the treatment conditions and organs except in the spleen and tumor after surgical resection of the brain tumor (**Supplemental Fig. 2**). Initial analysis of immunophenotypic changes in GL261-bearing mice revealed that the immune-suppressive granulocytic MDSCs (G-MDSCs) were proportionally increased in the blood of mice in response to dexamethasone treatment alone as well as mildly increased due to surgical resection (**Supplemental Figs. 3, 4**). Immune suppressive G-MDSCs were also increased in the recurrent tumors after resection and/or steroid treatment compared to mock resected animals (**Supplemental Fig. 3**). More strikingly, we observed a reduction in circulating T cells, including CD8+ T cells, following surgical resection alone, with a corresponding increase in CD8+ T cells in the bone marrow (**Fig. 2A-B**). These observations were repeated in the CT-2A resection model, and a similar decrease in CD8+ T cells in the blood was observed due to surgical resection, with a corresponding increase in CD8+ T cells in the bone marrow (**Fig. 2C-D, Supplemental Figs. 5-7**). These observations indicate that removing the tumor caused a reduction in circulating lymphocytes (**Fig. 2E**), and we next sought to determine whether tumor volume directly affected the levels of circulating T cells.

**Figure 2.**
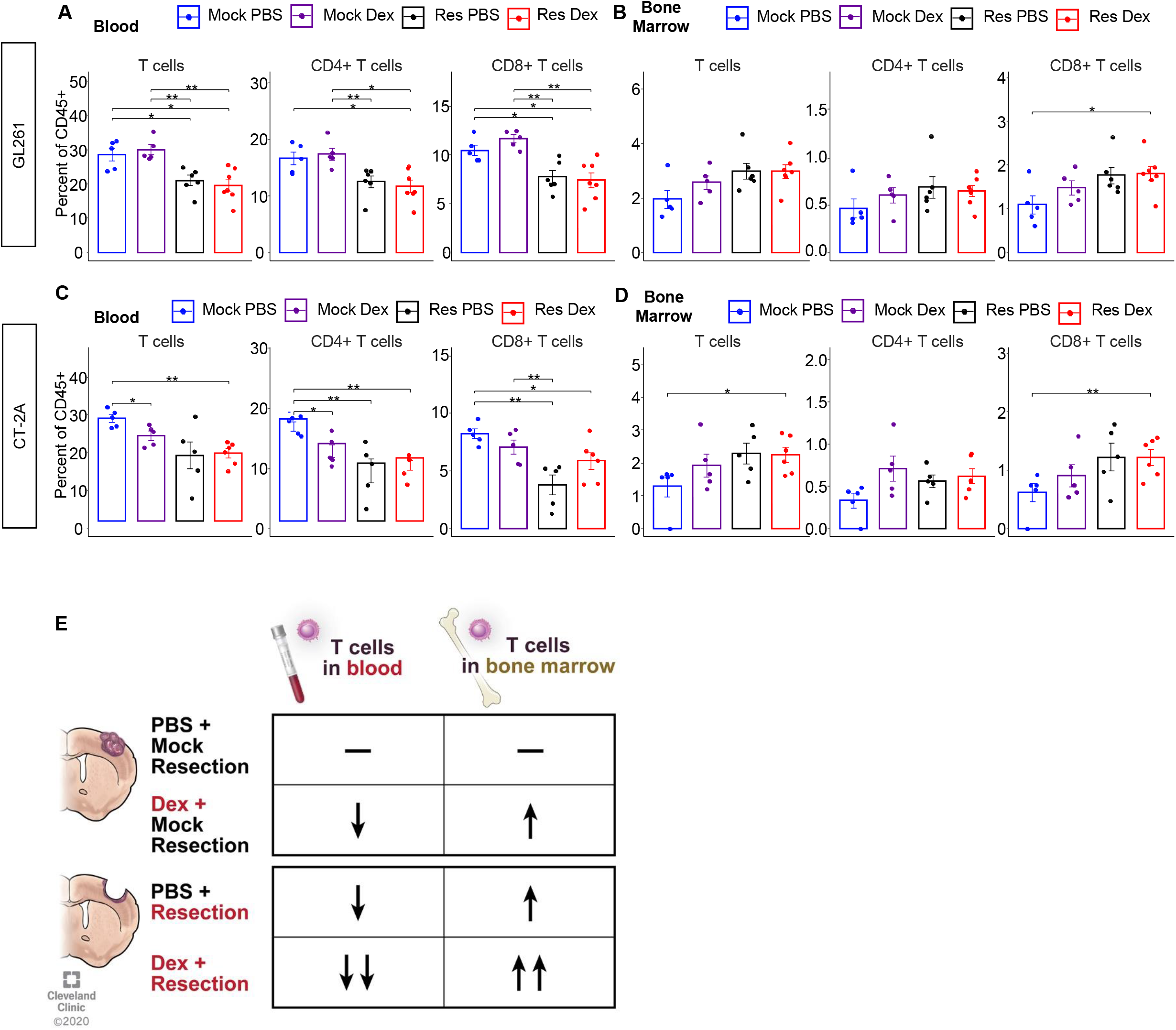
Surgical resection reduces T cells in the circulation and increases CD8+ T cells in the bone marrow. GL261-bearing mice were administered vehicle (n=7) or 4 μg dexamethasone daily (n=7) for 7 days starting on day 14 post-intracranial injection before being euthanized at day 21 for flow cytometry analysis of the blood and bone marrow (**A, B**). For the surgical resection model, mock PBS n=5, mock dexamethasone n=5, resection PBS n=6, and resection dexamethasone n=7. These studies were repeated using the CT-2A model of glioma, and T cell levels across treatment conditions were analyzed in the blood and bone marrow (**C, D**). Mock PBS n=5, mock dexamethasone n=5, resection PBS n=5, resection dexamethasone n=6. Summary graphic describing the gross overall T cell changes in the blood and bone marrow of each treatment (**E**). Student’s two-tailed t-test was performed for the comparisons in panels A, D, E; *p<0.05, **p<0.01, ***p<0.001.

### Murine tumor volume negatively correlates with circulating T cells but is masked by steroid treatment

To further understand how tumor volume and surgical resection might impact systemic immune parameters, we performed a volumetric analysis of the MRIs available at day 7 post-resection, the same day that flow cytometry was performed. Across a larger cohort of experimental mice, analysis of the bone marrow in the mock PBS, resection PBS, mock dexamethasone, and resection dexamethasone groups revealed a linear relationship between recurrent tumor volume and the quantity of T cells in the bone marrow (**Fig. 3A-D).** Of note, the number of CD8+ T cells was significantly higher in all groups compared to the combination of surgical resection and dexamethasone treatment, highlighting the combined impact of the two therapies. Further analysis of the blood lymphocyte counts revealed an inverse correlation between the number of T cells in the circulation and tumor volume (**Fig. 3E-H**). This inverse correlation was readily apparent when surgical resection was not performed, and similar trends were also present in the resection PBS group (**Fig. 3E-H**). In-depth analysis of recurrent tumor volume and its correlation with myeloid cell populations and lymphocytes within the spleen, blood, bone marrow, recurrent tumor and non-tumor cortex did not identify any other consistent correlations with recurrent tumor volume (**Supplemental Figs. 8-12**). Thus, our murine model suggests that T cells in the blood and marrow decrease and increase, respectively, with tumor volume, and that these relationships vanish when surgical resection and steroid treatment are both applied.

**Figure 3.**
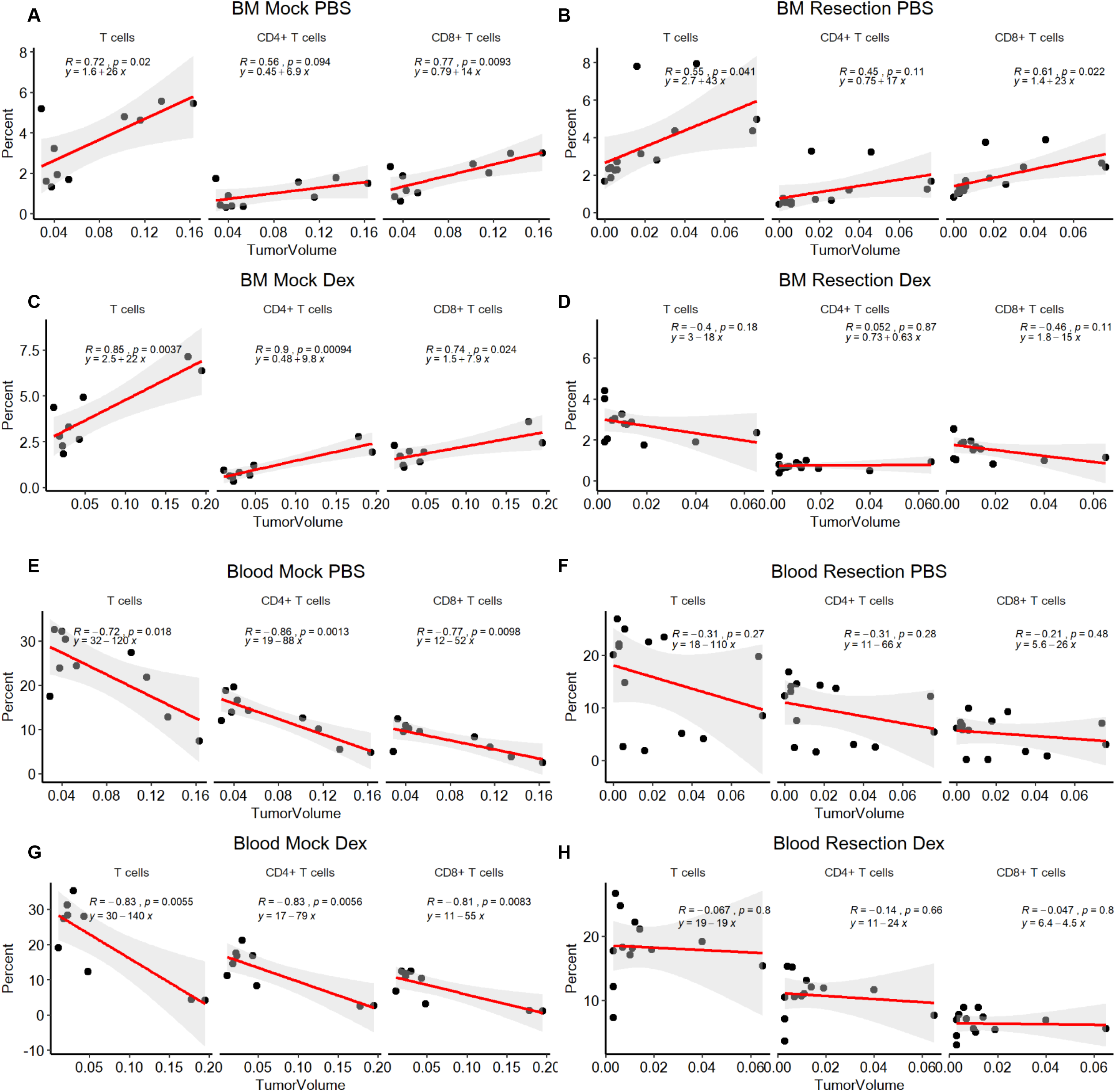
Tumor volume correlates with increased T cells in the bone marrow (BM) and reduced T cells in the blood. The bone marrow (**A-D**) and blood (**E-H**) are graphed as the percentage of T cells, CD4+ and CD8+ of total CD45+ cells, and the tumor volume. n=10 for mock PBS, n=9 for mock dexamethasone, n=14 for resection PBS, and n=13 for resection dexamethasone. Pearson correlation coefficients (R), p values and estimates of fitted line parameter are shown.

### Steroid-naïve GBM patients exhibit an inverse correlation between peripheral blood lymphocyte counts and progression-free and overall survival

To determine if relationships identified in our mouse model also existed in human patients with GBM, we assessed a clinical cohort of GBM patients who had an MRI as well as a complete blood count with differential (CBC w/ diff) prior to surgical resection. After analysis of patient records from a cohort of >400 GBM patients, we identified 95 GBM patients who met these criteria (**Table 1**). The demographics of the cohort utilized were analogous to the larger population of newly diagnosed GBM patients taken to surgery within our institution who did not meet the above criteria. Within the cohort of 95 newly diagnosed GBM patients, 61 patients had a CBC count prior to receiving any steroid treatment, while the other 34 patients had already received or were currently on steroids at the time of the CBC w/ diff, all prior to surgery. Univariate overall survival Kaplan-Meier analysis demonstrate that Stupp protocol, Tumor laterality, Resection type, Post operation MRI contrasted tumor volume, and age were all correlated with survival as expected (**Supplemental Fig. 13 A-F**). Analysis demonstrated that comparison of those patients who received steroids prior to surgery (n=34) and those who did not receive steroids prior to surgery (n=61) demonstrated the expected reduction in circulating lymphocytes, with no effect on the monocyte populations (**Fig. 4A, Supplemental Fig. 14A**). Additionally, we see that even with steroid treatment patients there is a trend toward surgical resection reducing peripheral blood lymphocyte counts, as compared to biopsy (**Supplemental Fig. 14B** Tumor volumes were then calculated from the MRIs corresponding to the time of CBC w/ diff, and we found that steroid-naïve patients demonstrated an inverse correlation between tumor volume and both lymphocyte percentage and absolute count by CBC w/ diff (**Fig. 4B, Supplemental Fig. 14C-D)**. In contrast, steroid-treated patients did not exhibit the same correlations (**Fig. 4C, Supplemental Fig. 14C-D)**. Univariate and multivariate Cox model analyses of the relationship between peripheral lymphocyte counts and progression-free survival (PFS) and overall survival (OS) were performed. In these analyses the PFS and OS were analyzed with the clinical variables: %Lymphocytes pre-surgery, Absolute Lymphocytes pre-surgery, Steroid treatment pre-surgery, Tumor Volume, Year Diagnosed, Age, Sex, Type of resection, Biopsy vs Resection, Tumor Laterality, KPS, 1p, 19q, ki67, and EGFR Amplification status. Within these analyses we have excluded the completion of Stupp protocol based on its dependence on other variables in the model such as type of resection, tumor laterality, biopsy vs resection, lymphocyte count, and residual tumor volume which are all already included in the models. Multivariate modeling was performed using these variables in a Cox Proportional-Hazards model using the R package glmulti with automated model selection set to identify the top 5 models based on Akaike’s Information Criterion (AIC), exhaustively searching the entire model space. Of these 5 models we display the top model for progression free survival (**Fig. 4D**) and Overall Survival (**Fig. 4E**), both of which include the absolute lymphocyte count at diagnosis; absolute lymphocyte count at diagnosis was also present in the four runner up models (**Supplemental Figures 15, 16),** which have similar AIC values and are thus close competitors. Importantly, other variables such as Biopsy vs resection, age, sex, 1p19q status, and tumor laterality are present in the progression free survival model and are known to affect GBM prognosis **(Fig. 4D, Supplemental Fig. 15A-D)**. Of note steroid treatment pre-op was also identified as a variable within the models of overall survival (**Fig. 4E, Supplemental Fig. 16A-D)**. After identifying that the absolute lymphocyte count is correlated with PFS and OS we sought to understand if the lymphocyte count is consistent over time in GBM patients based on their initial CBC/diff. In steroid-naïve patients, we identified that initial lymphocyte counts correlate with counts at recurrence (**Fig. 4F**) and in steroid naïve patients, with PFS and OS, but this is lost in steroid treated patients (**Supplemental Fig. 17A-B).**

**Table 1.** Patient Demographics table. Abbreviations: IDH: isocitrate dehydrogenase; MGMT: O-6-methylguanine-DNA methyl transferase; EGFR: epidermal growth factor receptor; STR: subtotal resection; NTR: near-total resection; GTR: gross-total resection; CBC: complete blood count; LITT: laser interstitial thermal therapy. Post-op CBC: n=40 in the no pre-op steroid group, n=27 in the yes pre-op steroid group. NS: p > 0.2

| Characteristic | Total Cohort | No Steroids Given<br>Preop Cohort | Steroids Given<br>Preop Cohort | Significant<br>Difference |
| --- | --- | --- | --- | --- |
|  | N (%) or Median Value (IQR) |  |  | p value |
| **Total Patients** | 95 | 61 (64.21%) | 34 (35.79%) |  |
| **Age at Diagnosis** | 63.72 (54.16-70.69) | 65.16 (55.25-72.65) | 59.45 (51.84-67.51) | NS |
| **Male Sex** | 68 (71.58%) | 41 (67.21%) | 27 (79.41%) | NS |
| **Karnofsky Performance Status** | 80 (80-90) | 80 (80-90) | 90 (75-90) | NS |
| **Ethnicity** |  |  |  |  |
| **Non-Hispanic White** | 86 (90.53%) | 56 (91.80%) | 30 (88.23%) | NS |
| **Laterality** |  |  |  | NS overall |
| **Left** | 41 (43.16%) | 25 (40.98%) | 16 (47.06%) |  |
| **Right** | 50 (52.63%) | 32 (52.46%) | 18 (52.94%) |  |
| **Bilateral** | 4 (4.21%) | 4 (6.56%) | 0 (0%) |  |
| **Lobe** |  |  |  | NS overall |
| **Deep** | 8 (8.42%) | 5 (8.20%) | 3 (8.82%) |  |
| **Multiple** | 18 (18.95%) | 10 (16.39%) | 8 (23.53%) |  |
| **Molecular Markers (if known of cohort)** |  |  |  |  |
| **1p Loss** | 11 (11.58%) | 6 (9.84%) | 5 (14.71%) | NS |
| **19q Loss** | 13 (13.68%) | 9 (14.75%) | 4 (11.76%) | NS |
| **IDH Mutated** | 2 (2.10%) | 1 (1.64%) | 1 (2.94%) | NS |
| **MGMT Methylated** | 13 (13.68%) | 7 (11.48%) | 6 (17.68%) | NS |
| **EGFR Amplification** | 40 (42.11%) | 26 (42.62%) | 14 (41.18%) | NS |
| **Ki-67** | 25% (15-50) | 27.5% (15.5-50) | 20% (15-50) | NS |
| **Initial Surgical Management** |  |  |  | NS overall |
| **Biopsy only** | 20 (21.05%) | 13 (21.31%) | 7 (20.59%) |  |
| **Laser Ablation** | 2 (2.11%) | 1 (1.64%) | 1 (2.94%) |  |
| **Surgical Resection** | 70 (73.68%) | 47 (77.05%) | 26 (76.47%) |  |
| **GTR** | 35 (36.84%) | 22 (36.06%) | 13 (38.24%) |  |
| **NTR** | 16 (16.67%) | 13 (21.31%) | 3 (8.82%) |  |
| **STR** | 19 (20.00%) | 12 (19.67%) | 7 (20.59%) |  |
| **Adjuvant Therapy** |  |  |  |  |
| **Stupp Protocol Completed** | 70 (73.68%) | 41 (67.21%) | 29 (85.29%) | 0.088 |
| **Volume of Contrast-Enhancing Tumor, preop (cc)** | 23.75 (14.01-41.77) | 24.20 (13.33-43.82) | 23.67 (15.67-40.57) | NS |
| **Resection only, preop (cc)** |  | 29.28 (14.12-48.94) | 24.96 (15.67-40.57) | NS |
| **Biopsy only, preop (cc)** |  | 17.69 (10.14-21.02) | 19.01 (17.08-44.51) | NS |
| **LITT only, preop (cc)** |  | 5.79 | 5.06 |  |
| **Volume of FLAIR Abnormality, preop (cc)** | 84.71 (44.61-127.78) | 76.94 (44.61-129.40) | 89.78 (45.70-124.26) | NS |
| **Preop Daily Dexamethasone Dose @ CBC (mg, if known)** | 0 (0-7) | 0 | 16 (12-16) | <0.0001 |
| **Resection only, preop, n=66** |  | 0 | 16 (12-16) |  |
| **Biopsy only, preop, n=20** |  | 0 | 12 (6-21) |  |
| **White Blood Cells, preop (x10<sup>3</sup>/μL), n=95** | 8.87 (6.44-11.84) | 7.57 (6.20-9.91) | 11.54 (8.82-14.85) | <0.0001 |
| **Percent Lymphocyte Count on CBC, preop, n=95** | 17.60 (8.40-25.70) | 21.40 (17.40-28.95) | 7.15 (5.43-11.28) | <0.0001 |
| **Resection only, preop, n=73** |  | 21.3 (17.2-28.8) | 7.1 (5.08-11.28) | <0.0001 |
| **Biopsy only, preop, n=20** |  | 25.7 (17.6-34.3) | 8.2 (6.3-21.8) | 0.009 |
| **Absolute Lymphocyte Count on CBC, preop (x10<sup>3</sup>/μL), n=95** | 1.42 (0.90-2.24) | 1.73 (1.22-2.37) | 0.88 (0.53-1.21) | 0.0002 |
| **Resection only, preop** |  | 1.95 (1.20-2.37) | 0.875 (0.49-1.14) | 0.001 |
| **Biopsy only, preop** |  | 1.73 (1.19-2.53) | 1.23 (0.64-2.78) | NS |
| **Percent Monocyte Count on CBC, preop, n=95** | 6.0 (3.7-8.0) | 6.9 (5.3-8.1) | 5.8 (1.8-6.7) | 0.014 |
| **Absolute Monocyte Count on CBC, preop (x10<sup>3</sup>/μL), n=95** | 0.52 (0.34-0.71) | 0.52 (0.39-0.68) | 0.50 (0.15-0.85) | NS |
| **Volume of Contrast-Enhancing Tumor, post-op (cc, including biopsies, not including LITT, if post-op CBC obtained)** | 0.54 (0-7.95) | 0.83 (0-7.73) | 0.35 (0-13.70) | NS |
| **Resection only, post-op, n=29** |  | 0.50 (0-4.51) | 0 (0-0.54) |  |
| **Biopsy only, post-op, n=5** |  | 11.98 (8.82-18.76) | 18.93 (15.95-45.03) |  |
| **Post-op Daily Dexamethasone Dose @ CBC (mg, if known)** | 16 (8-16) | 16 (8-16) | 16 (8-16) | NS |
| **Resection only, post-op, n=55** |  | 16 (8-16) | 16 (12-16) | NS |
| **Biopsy only, post-op, n=10** |  | 8 (5-14) | 12 (7-22) | NS |
| **White Blood Cells, post-op (x10<sup>3</sup>/μL), n=67** | 13.27 (9.03-16.15) | 13.36 (8.42-16.03) | 12.75 (9.62-16.35) | NS |
| **Percent Lymphocyte Count on CBC, post-op, n=67** | 8.2 (4.8-14.1) | 8.45 (5.43-14.1) | 6.5 (4.7-14.4) | NS |
| **Resection only, post-op, n=55** |  | 8.7 (5.5-14.1) | 6.25 (4.35-11.58) |  |
| **Biopsy only, post-op, n=10** |  | 8.0 (4.5-14.6) | 12.85 (9.83-18.30) |  |
| **Absolute Lymphocyte Count on CBC, post-op (x10<sup>3</sup>/μL), n=67** | 0.96 (0.67-1.54) | 0.96 (0.66-1.59) | 0.96 (0.68-1.42) | NS |
| **Resection only, post-op, n=55** |  | 1.00 (0.67-1.61) | 0.74 (0.65-1.22) |  |
| **Biopsy only, post-op, n=10** |  | 0.86 (0.53-1.66) | 1.34 (0.93-1.76) |  |
| **Percent Monocyte Count on CBC, post-op, n=67** | 6.8 (4.8-9.0) | 6.1 (3.13-8.5) | 7.6 (5.2-9.4) | 0.038 |
| **Absolute Monocyte Count on CBC, post-op (x10<sup>3</sup>/μL), n=67** | 0.88 (0.52-1.11) | 0.72 (0.44-1.06) | 1.01 (0.82-1.16) | 0.042 |
| **Progression-Free Survival (mo)** | 5.56 (3.02-11.31) | 5.03 (2.86-8.52) | 8.38 (3.48-13.28) | 0.125 |
| **Overall Survival (mo)** | 11.23 (4.87-20.9) | 10.4 (4.73-17.67) | 17.17 (6.67-29.17) | 0.085 |

Post op CBC: n=40 in no preop steroid group, n=27 in yes preop steroid group
NS: p > 0.2

**Figure 4.**
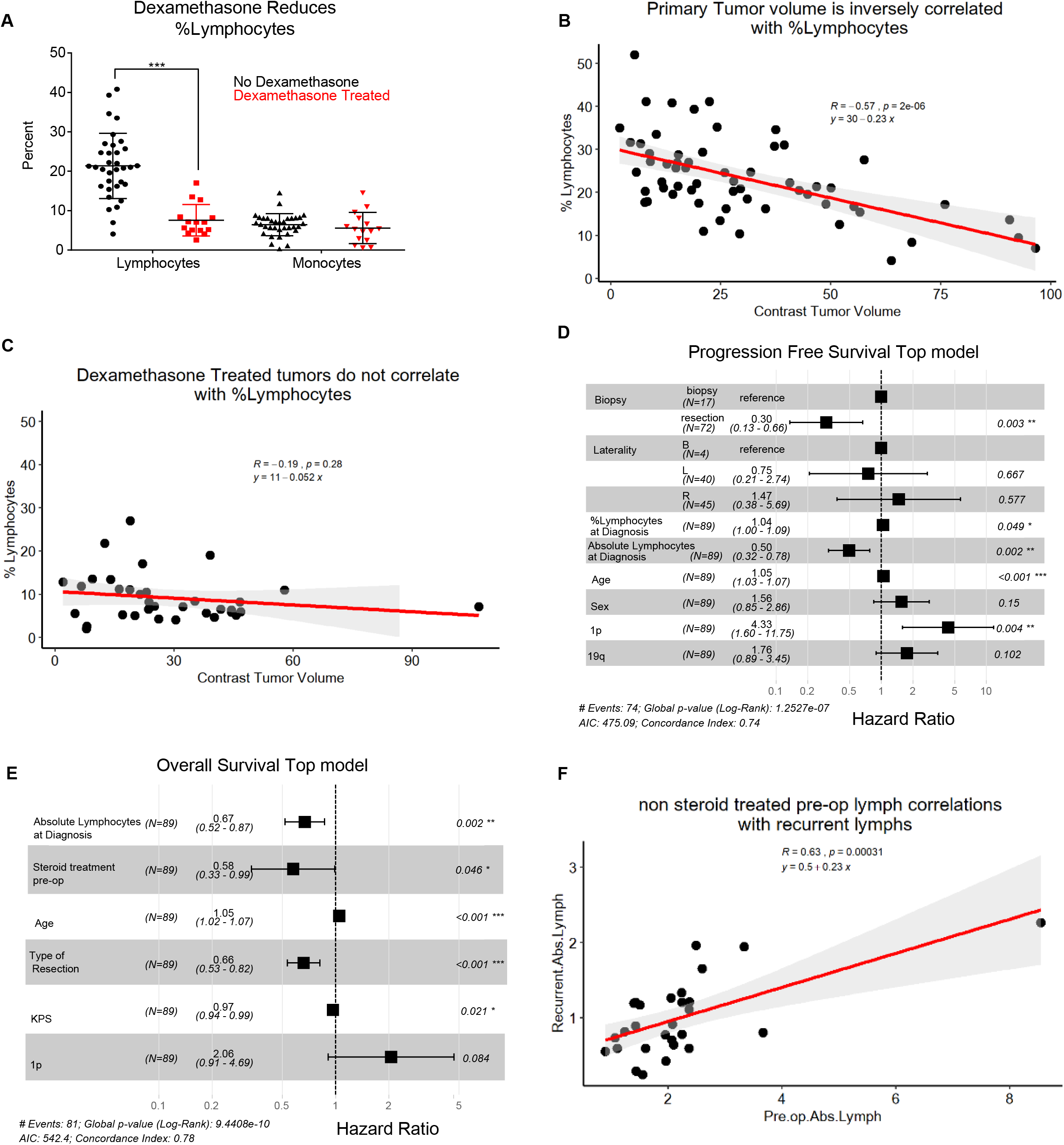
Patient cohort validates that tumor volume negatively correlates with lymphocytes prior to surgery and steroid treatment. Using a cohort of n=95 patients prior to surgical resection, the first CBC with differential was used to compare % lymphocytes between dexamethasone-treated- and non-dexamethasone-treated patients (**A**). Percentages of lymphocytes vs tumor volumes (assessed by MRI) of patients without dexamethasone treatment (**B**). Percentages of lymphocytes vs tumor volumes for those patients treated with dexamethasone prior to surgery who also had a matching MRI at the time of CBC w/ diff (**C**). Multivariate Cox proportional hazards model of Progression Free Survival (**D**). Multivariable Cox proportional hazards model of Overall survival (**E**). Absolute lymphocyte counts at recurrence and pre-op are correlated for patients not treated with steroids at the time of pre op absolute lymphocyte count (**F**). The correlation coefficient (R), p value and fitted line parameter are shown.

## Discussion

As immunotherapies for GBM evolve, the context in which they will be administered, and more specifically the impact of existing standard-of-care paradigms, must be considered. In particular, interactions with surgical resection, steroids, chemotherapy, and radiation, must be assessed, as must the importance of their timings relative to administration of immune-modulating agents. Recently, immunotherapy in GBM failed in the CheckMate 143 clinical trial^17^, where recurrent GBM patients who had previously failed standard-of-care therapy were evaluated for their response to anti-PD1 immune checkpoint therapy. Following this trial, a smaller study was initiated with therapy delivered prior to surgical resection (neoadjuvant)^14^, and moderate but promising effects were observed. New clinical trials testing neoadjuvant immunotherapy are increasingly being attempted. Preclinical modeling of these paradigms is critical, as it will generate hypotheses regarding how surgery, steroids, and the immune system interact.

Using mouse GBM models, we determined that surgical resection alone leads to a reduction in circulating T cells, in addition to the known impact of steroid treatment^18–20^, which is also being revisited in terms of its clinical use. Many groups have shown that T cell abundance is one of multiple factors influencing immunotherapy response. Our murine mock-resection model enables studies of cohorts of mice with similar time of surgery, time under anesthesia, blood loss, dural penetration, and disruption of native cortex. This allowed us to conclude that regardless of the initial factor causing a reduction in lymphocytes, the simple act of tumor debulking also causes a reduction of T cells in the circulation. Likewise, in a cohort of GBM patients, we see the same trend toward surgical resection reducing peripheral blood lymphocyte counts, as compared to biopsy, even when steroids are present (**Supplemental Fig. 14B**). Furthermore, T cell sequestration in the bone marrow in the setting of newly diagnosed intracranial tumors has been previously described^3^. We observe the same accumulation of T cells in the bone marrow in murine models of surgically resected, recurrent tumors and additionally that this accumulation varies with tumor volume. As mice studied are syngeneic, decreasing lymphocyte counts is unlikely to be due to differences in host immune constitution. We also observe an inverse relation between peripheral blood lymphocyte counts and tumor volume in steroid-naïve human patients. In these patients decreased pre-operative lymphocyte counts correlated with worse progression-free- and overall survival in multivariate models. Additionally, those patients who had steroid treatment prior to surgical resection at the time of the CBC w/diff had a worse overall survival, which could be due to the severity of disease but could also be reducing the ability to respond to the tumor.

Critically, contrasted pre-operative tumor volume not being a significant risk factor in uni- and multi-variate models of progression-free- and overall survival suggests that mechanistically, preoperative lymphocyte counts are not merely surrogate measures of tumor volume/stage but instead may reflect a patient’s underlying ability to mount an immune response to GBM. Similarly, steroid use dropping out of significance in multivariate hazard ratio modeling progression free survival suggests that a patient’s intrinsic or at least pre-operative immune status is a stronger determinant of the risk of disease progression.

Peripheral lymphocyte counts at diagnosis, prior to steroid administration, may reveal which patients are more likely to benefit from immuno-therapies. Patients with higher baseline lymphocyte counts might be more likely to benefit from neoadjuvant checkpoint blockade and patients with lower counts might benefit from receiving T cell-activating or -mobilizing therapies. A more complete understanding of a patient’s baseline immune status and the temporal effects of administration of therapies on the immune system is needed to enable separations of patients into subgroups more likely to benefit from a given immunotherapy. Developing such initial assessments will likely be critical to the success of GBM immuno-therapies in clinical trials.

## Methods

### Syngenic Tumor Resection Models

GL261 mice were acquired from the NCI and CT-2A mice were obtained from Dr. Thomas Seyfried (Boston University). Six-week-old aged-matched male C57BL/6J mice from Jackson Labs (000664) were anesthetized using isoflurane and then intracranially injected into the left cerebral hemisphere with 20,000 GL261 or CT-2A cells in 5 μl of RPMI medium using a stereotactic frame. This model has been established in our laboratory using neurological symptoms as an indicating endpoint; median survival times are approximately 20 days^21^.

On the day of surgical resection, 7 days after intracranial implantation of glioma cell lines (GL261, CT-2A), mice were taken to the MRI suite, and successful tumor implantation was confirmed by T2-weighted volumetric brain scans of the animals. After completion of the MRI scan, the animals were taken into the surgical procedure suite. Throughout the entire procedure, sterile aseptic techniques were used. Mice were anesthetized with 1.5%-5% isoflurane and 100% oxygen using an anesthesia vaporizer and monitored. Upon loss of hind-foot withdrawal and corneal reflex to gentle touch with a cotton swab, the mice were deemed ready for surgery. The fur along the left aspect of the cranium was shaved, and the mice were placed in a stereotactic frame with an adapted nose cone for continuous inhaled anesthetic delivery throughout the procedure. The cranium and future incision site were cleaned with 10% povidone-iodine (Medline, 53329-945-09) x3, the region around the surgical site was draped, and the operating microscope was brought into the field (Leica Microsystems, 6x scope). Subcutaneous lidocaine was injected along the planned incision. A #15 blade scalpel (Futura, SMS215) was used to make a linear cranial-caudal incision through the skin to the cranium, and the skin flaps were retracted using 5-0 polypropylene sutures (Ethicon). The periosteum was scraped using a #4 Pennfield retractor. Using an electric handheld drill with a 1.6 mm carbide round bur (Roboz Surgical Instruments, RS-6280C-5), copious irrigation with sterile saline, and suctioning, a 4 mm craniotomy was generated overlying the point of intracranial injection. Bony fragments were removed using micro forceps. The dura was incised using micro-scissors, and hemostasis was achieved by copious irrigation and oxidized regenerated cellulose matrix (Surgicel). A corticectomy was performed using a medium #5 Fukushima suction tip and tissue retraction with #4 Pennfield retractor. The gross tumor was visualized, and using a combination of suction, retraction, and microdissection, the tumor was debulked until no visible gross tumor remained. The surgical cavity was irrigated and inspected for visible tumor. Upon satisfactory visualization of only normal brain, the surgical cavity was lined with oxidized regenerated cellulose matrix (Surgicel). A larger piece of cellulose matrix was placed over the overlying dura. The retracting sutures were removed, and the skin was approximated using a running simple stitch using 6-0 polypropylene suture (Ethicon). The microscope was removed from the field, and the mouse was taken out of the stereotactic frame. Sterile normal saline (1 mL) was injected subcutaneously for hydration, and one dose of 0.1 mg/kg buprenorphine was given for analgesia before recovery. The mice were treated with either PBS or dexamethasone (Sigma-Aldrich, 4 μg) intraperitoneally. The mice were then placed on a heating pad and allowed to emerge from anesthesia. A second dose of buprenorphine was given 4-6 hours later on the day of surgery and another the following morning. After the third dose, buprenorphine was given up to three times daily PRN if the animals appeared to be in pain. Mice received 1 ml sterile normal saline daily for 4 days post operatively to prevent dehydration. Animals were provided with prophylactic antibiotics (neomycin, 500 μg/ml) and analgesics (ibuprofen, 200 μg/ml) added to their drinking water. Dexamethasone (Sigma-Aldrich, 4 μg) or PBS was administered intraperitoneally daily in the afternoon for the duration of the experiments.

### MRI

Mice were imaged at 7 days post-intracranial implantation and 24 h and 14 days post-tumor resection. MRI acquisitions were carried out on a 7 T MRI scanner (Bruker BioSpec 70/20 USR; Billerica, MA) using a 23-mm volume coil setup. Animals were anesthetized with an isoflurane/oxygen mixture (1-3%, VetFlo System, Kent Scientific) throughout the scan acquisition, with respiration and body temperature monitored via a physiological monitoring system (SA Instruments, Stoney Brook, NY). The animal’s head was consistently positioned within the 23-mm volume coil to ensure that the entire brain region was being scanned and that slice-to-slice comparisons could be made between the two time points. The imaging protocol utilized was a multi-slice acquisition (axial) fast-spin echo (FSE) sequence (T2-TURBORARE, Paravision 6.0) to provide structural information of the tumor volume and location. The T2-weighted FSE sequence was run with the following parameters: FOV = 1.8 × 1.8 cm, slice thickness of 0.5 mm, matrix size 180 × 180, TE = 50 ms (Echo Spacing = 7.0 and ETL = 16), TR = 4550 ms, and SA=6 with TA = 5 min per animal.

### Edema Scoring

From the volumetric T2-weighted axial MRI slices obtained for each mouse, the overall gross tumor was identified and highlighted using BrainLab 3.0 software. The visible T2 hyper-intensities outside of the tumor volume were identified as the surrounding vasogenic edema and scored on a scale of 0-3 to correlate tumor volume with edema volume. Importantly, the individual analyzing each MRI image was blinded to the treatment the mice received until after the edema was analyzed for all murine subjects.

### Patient Sample Volumetric

BrainLab 3.0 software was utilized to analyze the axial MRI slices of each mouse. The overall tumor area was outlined manually for each image in a treatment-blinded manner.

### Flow Cytometry

Antibody staining and flow cytometry were performed as previously described by our laboratory^22, 23^. Briefly, at the designated time point, 1 week after surgical resection, blood was collected in EDTA tubes form terminal bleeds, and the spleen, bone marrow, tumor and non-tumor tissue harvested and mechanically dissociated using 40 μM cell strainers. Each sample was then stained for live/dead cells using a Fixable Blue Dead Cell Stain Kit (Thermo Fisher Scientific, Catalog # L23105) and blocked using Fc Receptor block (Miltenyi Biotec 130-092-575). Next, samples were split into two parts for myeloid and lymphoid panel staining. The myeloid panel included: live/dead UV, CD45, CD11b, CD11C, IA/E, CD103, Ly6G, Ly6C, CD68, and Ki67, and the lymphoid panel included: live/dead UV, CD45, CD3, CD4, CD8, PD1, NK1.1, CD25, CD69, and FoxP3. Antibodies were obtained from Biolegend (San Diego, CA) for analysis of mouse immune profile and included fluorophore-conjugated anti-Ly6C (Clone HK1.4, Catalog # 128024), anti-Ly6G (Clone A8, Catalog # 127618), anti-CD11b (Clone M1/70, Catalog # 101212), anti-CD68 (Clone FA-11, Catalog # 137024), anti-I-A/I-E (Clone M5/114.15.2, Catalog # 107606), anti-CD11c (Clone N418, Catalog # 117330), anti-CD3 (Clone 145-2C11, Catalog # 100330), anti-CD4 (Clone GK1.5, Catalog # 100422), anti-CD8 (Clone 53-6.7, Catalog # 100712), anti-NK1.1 (Clone PK136, Catalog # 108741), anti-CD45 (Clone 30-F11, Catalog # 103132), anti-Ki-67 (Clone 16A8, Catalog # 652404). Gating for MDSCs was performed using FlowJo v10, and M-MDSCs were identified by (Singlets/Live/CD45+/CD11b+/CD68-/IAIE-/Ly6G-/LyC+) and G-MDSCs by (Singlets/Live/CD45+/CD11b+/CD68-/IAIE-/Ly6C-/Ly6G+). T cells were identified by (Singlets/Live/CD45+/CD3+/NK1.1-). CD4+ T cells were identified by (Singlets/Live/CD45+/CD3+/NK1.1-/CD4+/CD8-), CD8+ T cells by (Singlets/Live/CD45+/CD3+/NK1.1-/CD4-/CD8+), NK cells by (Singlets/Live/CD45+/NK1.1+), T regulatory cells by (Singlets/Live/CD45+/CD3+/CD4+/FoxP3+), and macrophages by (Singlets/Live/CD45+/CD11b+/CD68+/IAIE+). CD45+ cells are graphed as a percentage of live cells, while all other populations are graphed as percentage of live/single/CD45+ cells.

### Statistics

GraphPad Prism and R (version 4.0.2) were used. Times to events (progression or death) were modeled using Cox proportional hazards models. Kaplan Meier survival curve differences were assessed by log-rank tests. The R package survival was used to compute log rank tests and to fit Cox models in R; the R package survminer was used to form Kapan-Meier plots and forest plots of hazard ratio estimates. Automated exhaustive Cox model space searches were performed using the R package glmulti with Akaike’s Information Criterion (AIC) as the model selection criterion. Relative to the Bayesian Information Criterion (BIC), AIC tends to select larger models; both metrics strike balances between desires to increase goodness-of-fit and desires to fit fewer model parameters. Other data (e.g. flow cytometry data) was analyzed using the R packages ggplot2 and ggpubr, which integrate basic statistical tests with plotting. P-values were considered statistically significant at *p<0.05, **p<0.01, and ***p<0.001.

## Acknowledgments

We thank the members of the Lathia laboratory for insightful discussion and constructive comments on the manuscript. We thank the Lerner Research Institute Flow Cytometry Core for their assistance. We thank Amanda Mendelsohn and the Center for Medical Art and Photography at the Cleveland Clinic for providing illustrations.

## Author Contributions

BO, TA, MG, MV, PF, and JL provided conceptualization and design. BO, TA, DB, AR, SJ, CA, performed the experiments. BO, TA, MG, DB, AR CA, TR, AM, MA, MV, PF, JL analyzed the data. BO, TA, MG, EM, TR, MV, PF, JL wrote the manuscript. BO, JL provided financial support. All authors provided final approval of the manuscript.

## Supplemental Figure Legends

**Supplemental Figure 1.**
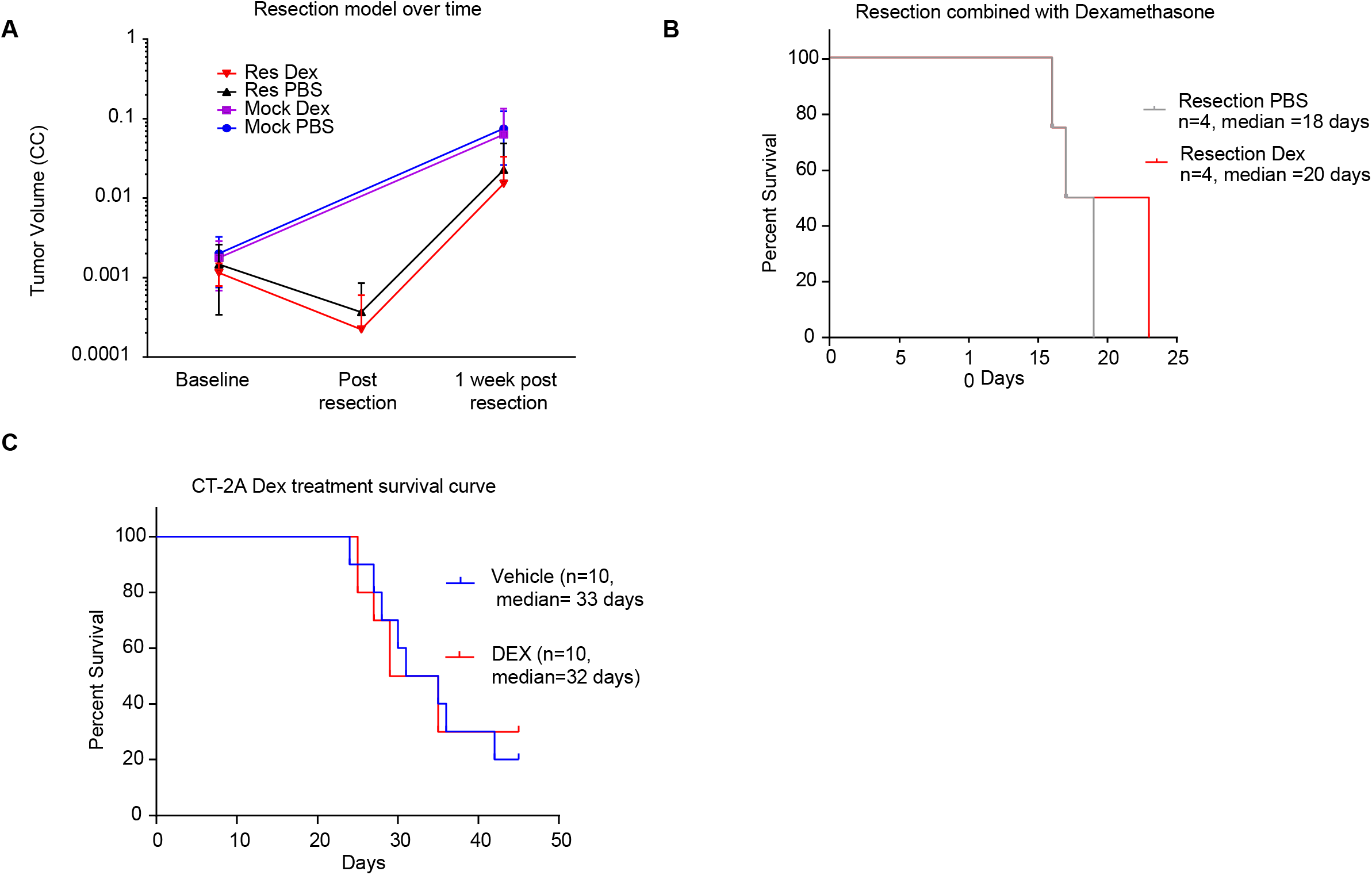
Mouse model of resection (n=7 per group) comparing the tumor volumes at baseline, post-resection, and 1 week post-resection. Note that the mock resected mice from this cohort did not receive back-to-back MRI on the same day as was performed for the resection cohort (**A**). Survival analysis of resection PBS-vs resection dexamethasone-treated animals (n=4 per group) (**B**). CT-2A-bearing vehicle- and dexamethasone-treated mice without surgical resection (n=10 mice per group) demonstrated no difference in survival, with median survival values of 33 and 32 days, respectively (**C**). Survival curve analysis was performed in GraphPad Prism using log-rank tests to obtain p values.

**Supplemental Figure 2.**
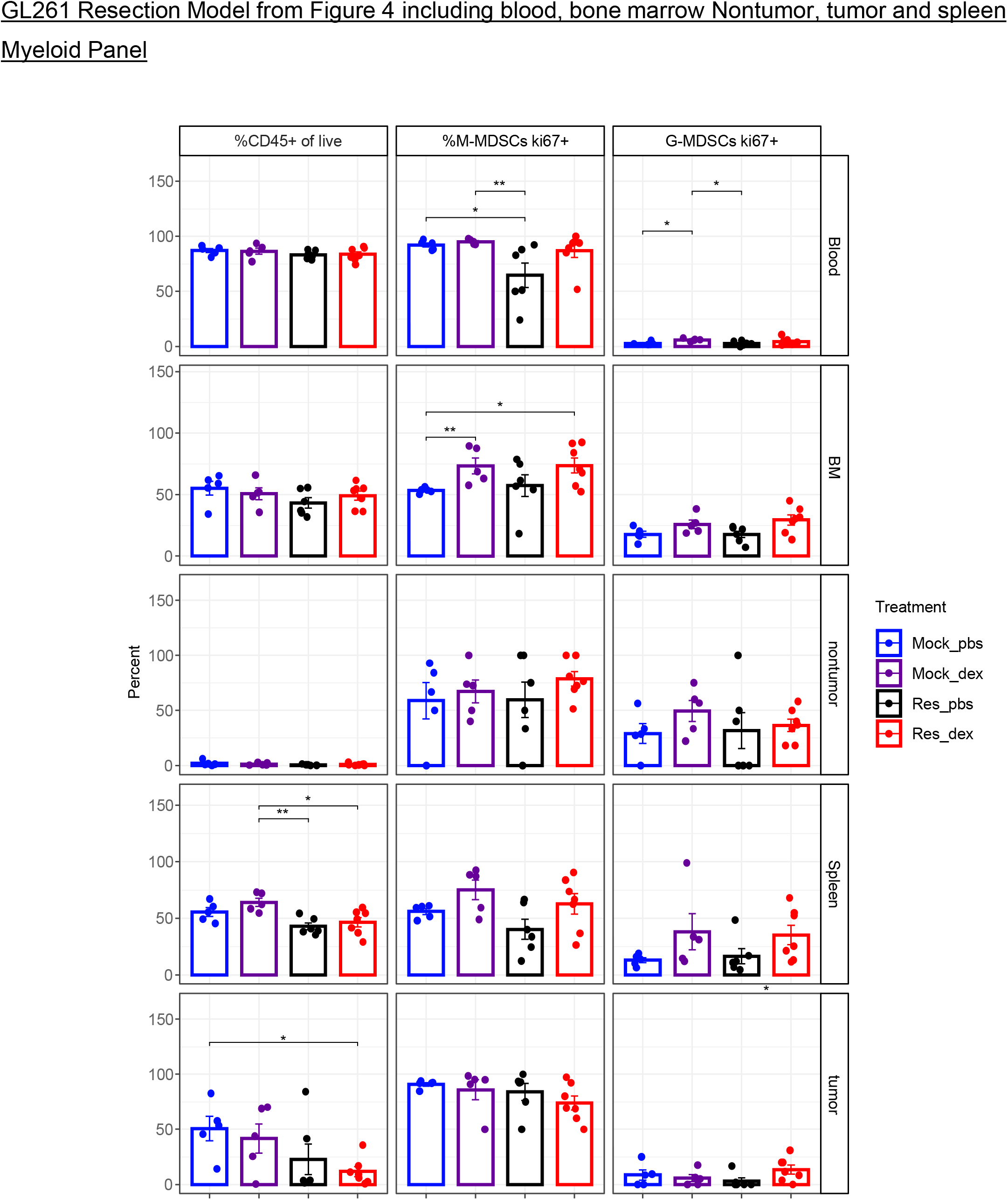
GL261-bearing mice as described in Figure 1 and Figure 2, including the groups mock PBS, mock dexamethasone, resection PBS, and resection dexamethasone, were evaluated via flow cytometry for % CD45+ cells of live cells, % Ki67+ M-MDSCs, and % Ki67+ G-MDSCs in the blood, bone marrow, non-tumor cortex, spleen, and tumor. Student’s two-tailed t-tests were used to perform the comparisons; *p<0.05, **p<0.01, ***p<0.001.

**Supplemental Figure 3.**
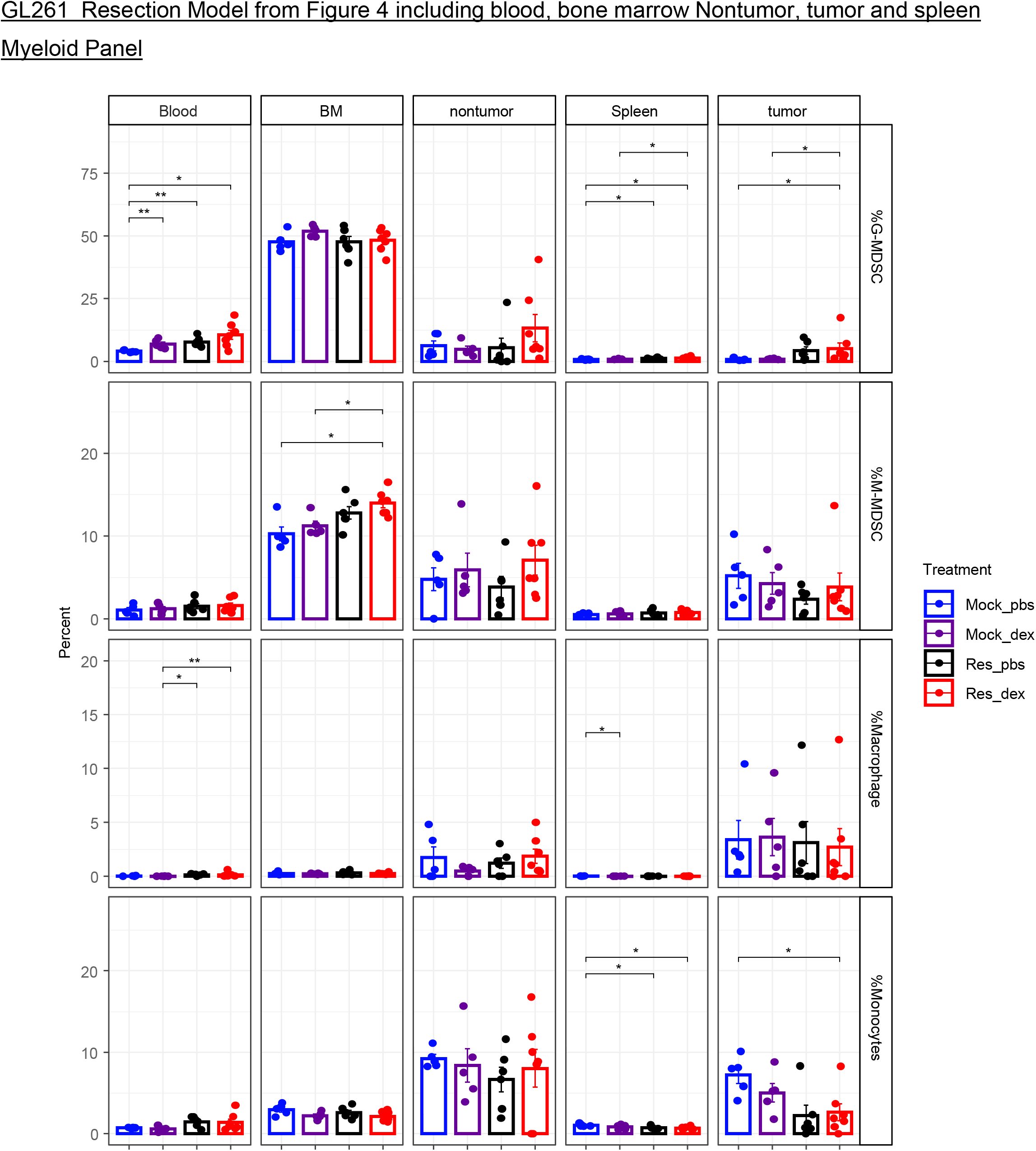
GL261-bearing mice as described in Figure 1 and Figure 2, including the groups mock PBS, mock dexamethasone, resection PBS, and resection dexamethasone were evaluated via flow cytometry for G-MDSCs, M-MDSCs, macrophages, and monocytes in the blood, bone marrow, non-tumor cortex, spleen, and tumor. Note: Blood and bone marrow G-MDSCs and M-MDSCs are shown in Figure 2C, D but are also shown globally with other organs for comparisons. T-tests were used to compare groups; *p<0.05, **p<0.01, ***p<0.001.

**Supplemental Figure 4.**
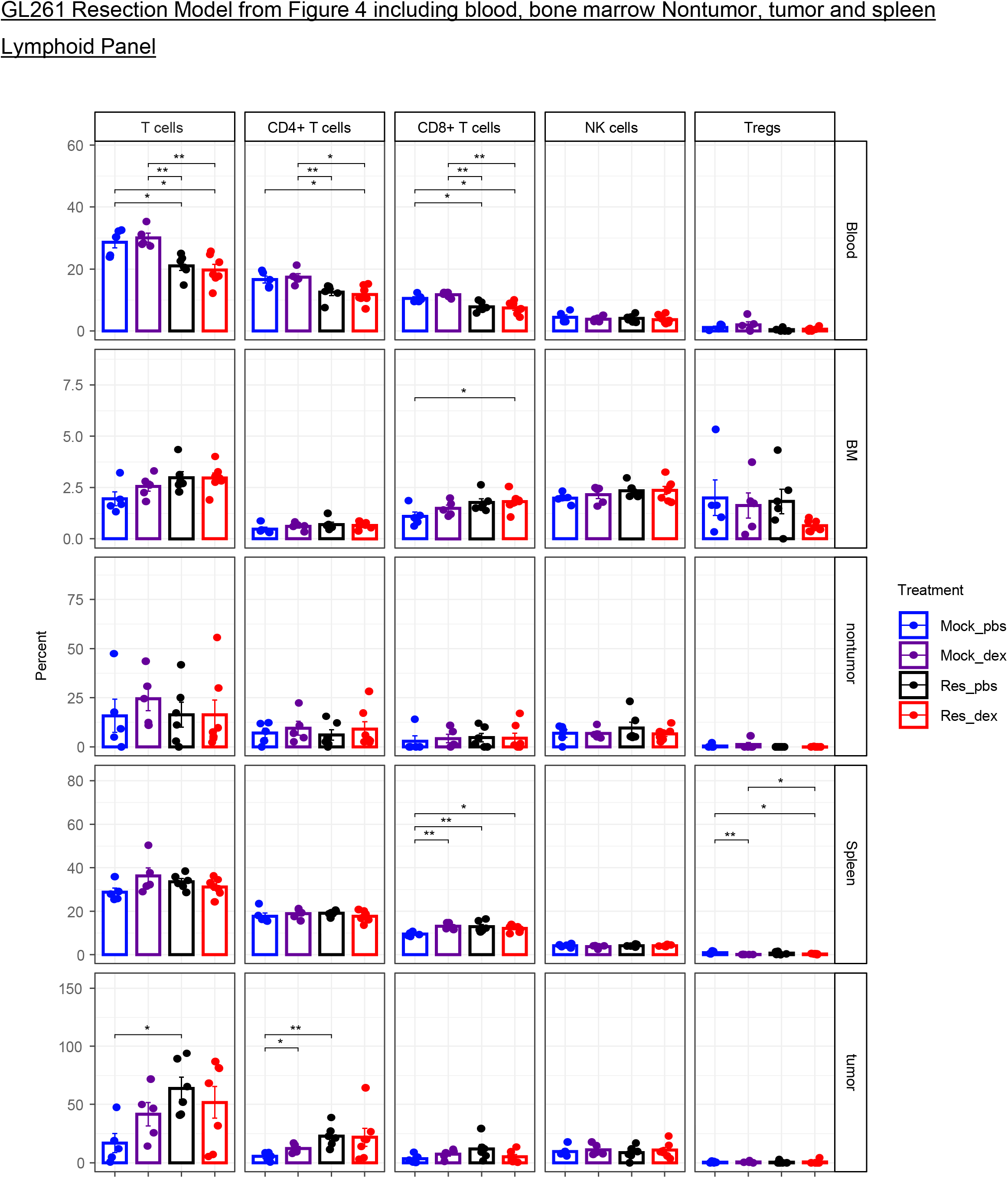
GL261-bearing mice as described in Figure 1 and Figure 2, including the groups mock PBS, mock dexamethasone, resection PBS, and resection dexamethasone, were evaluated via flow cytometry for T cells, CD4+ T cells, CD8+ T cells, NK cells, and T-regulatory cells in the blood, bone marrow, non-tumor cortex, spleen, and tumor. Note: Blood and bone marrow T cell populations are shown in Figure 2E, F but are shown globally with other organs here for comparison. Groups were compared by t-tests; *p<0.05, **p<0.01, ***p<0.001.

**Supplemental Figure 5.**
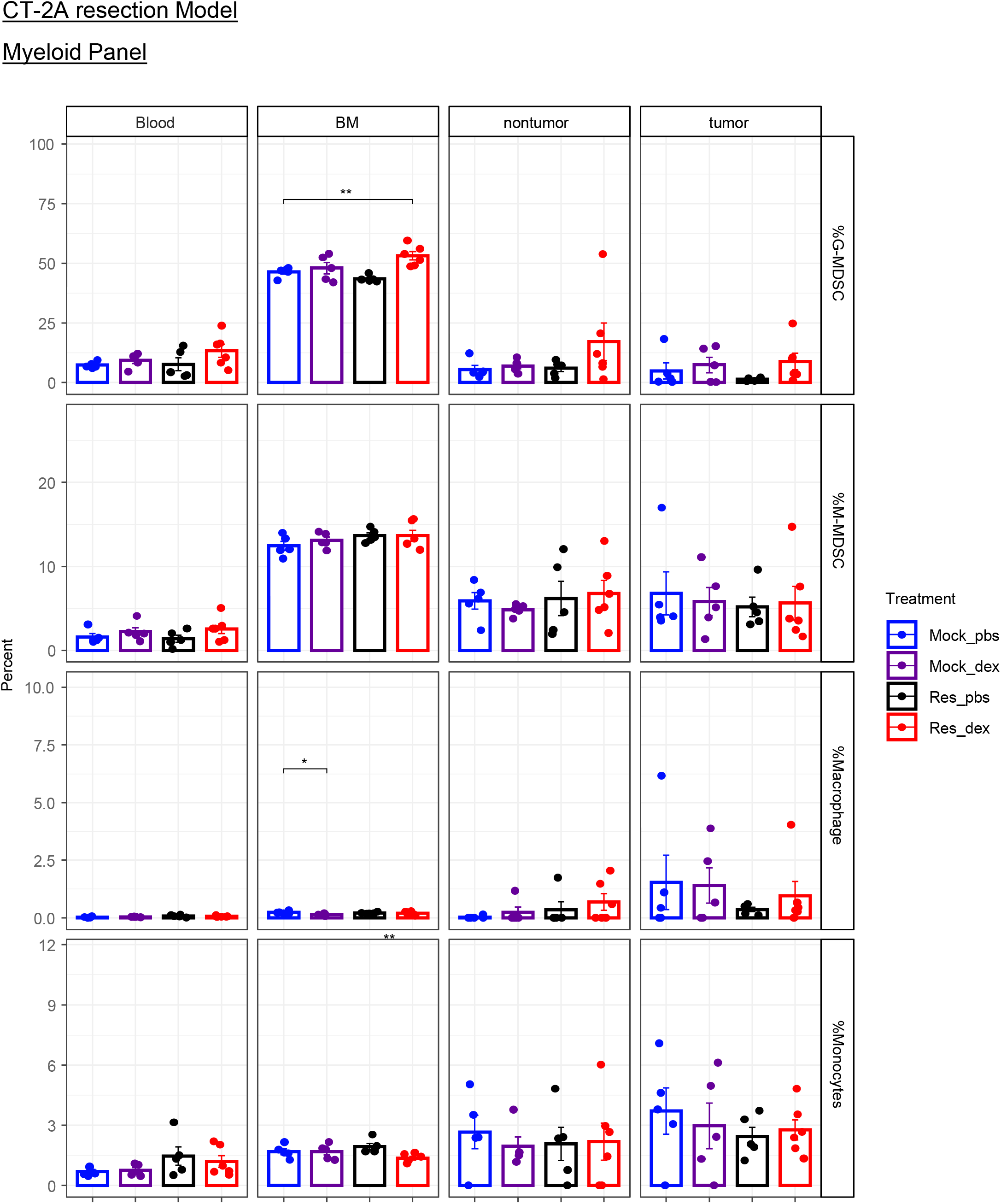
CT-2A-bearing mice as described in Figure 1 and Figure 2, including the groups mock PBS, mock dexamethasone, resection PBS, and resection dexamethasone, were evaluated via flow cytometry for G-MDSCs, M-MDSCs, macrophages, and monocytes in the blood, bone marrow, non-tumor cortex, spleen, and tumor. Groups were compared by t-tests; *p<0.05, **p<0.01, ***p<0.001.

**Supplemental Figure 6.**
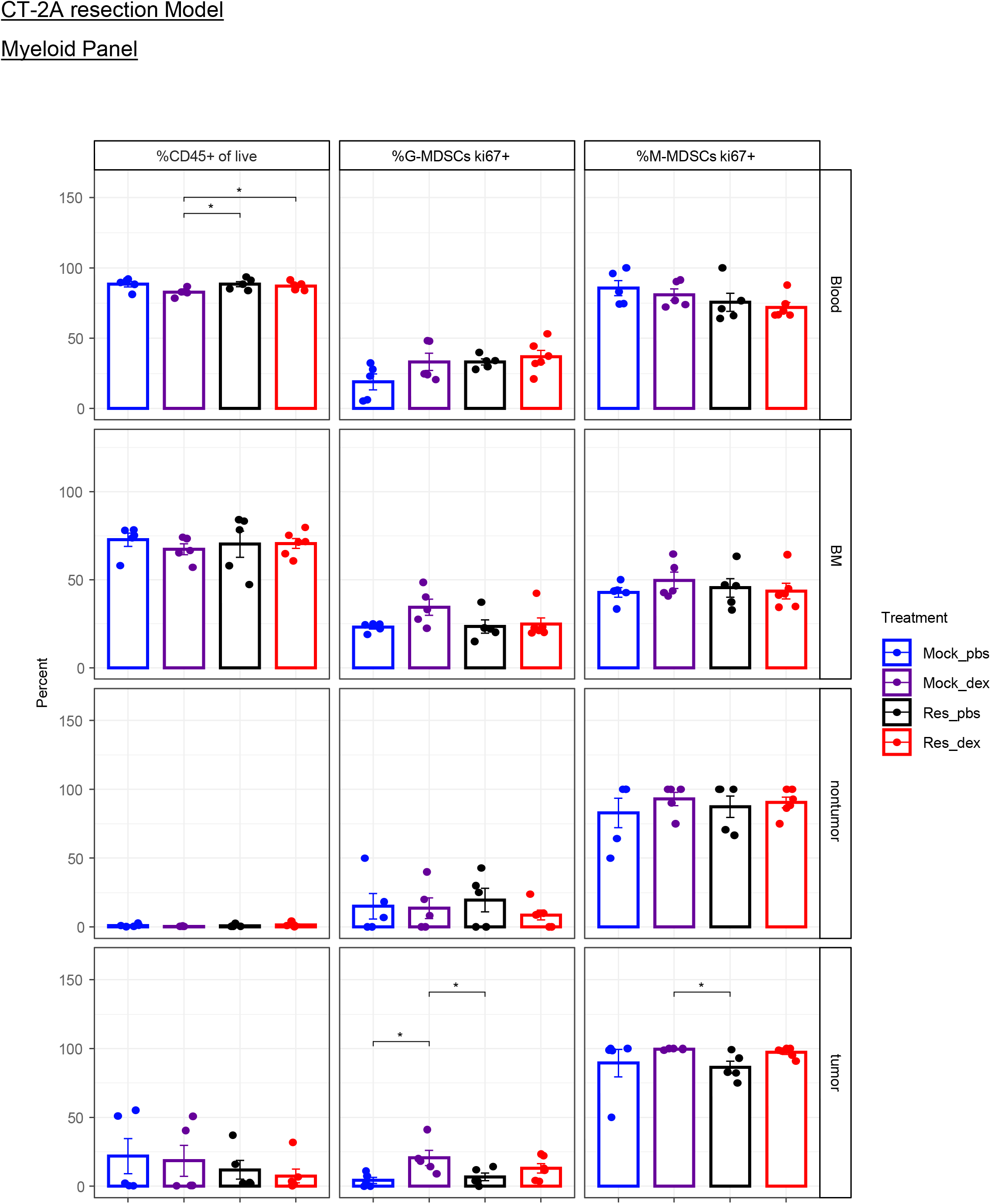
CT-2A-bearing mice as described in Figure 1 and Figure 2, including the groups mock PBS, mock dexamethasone, resection PBS, and resection dexamethasone, were evaluated via flow cytometry for % CD45+ cells of live cells, % Ki67+ M-MDSCs, and % Ki67+ G-MDSCs in the blood, bone marrow, non-tumor cortex, spleen, and tumor. Groups were compared by t-tests; *p<0.05, **p<0.01, ***p<0.001.

**Supplemental Figure 7.**
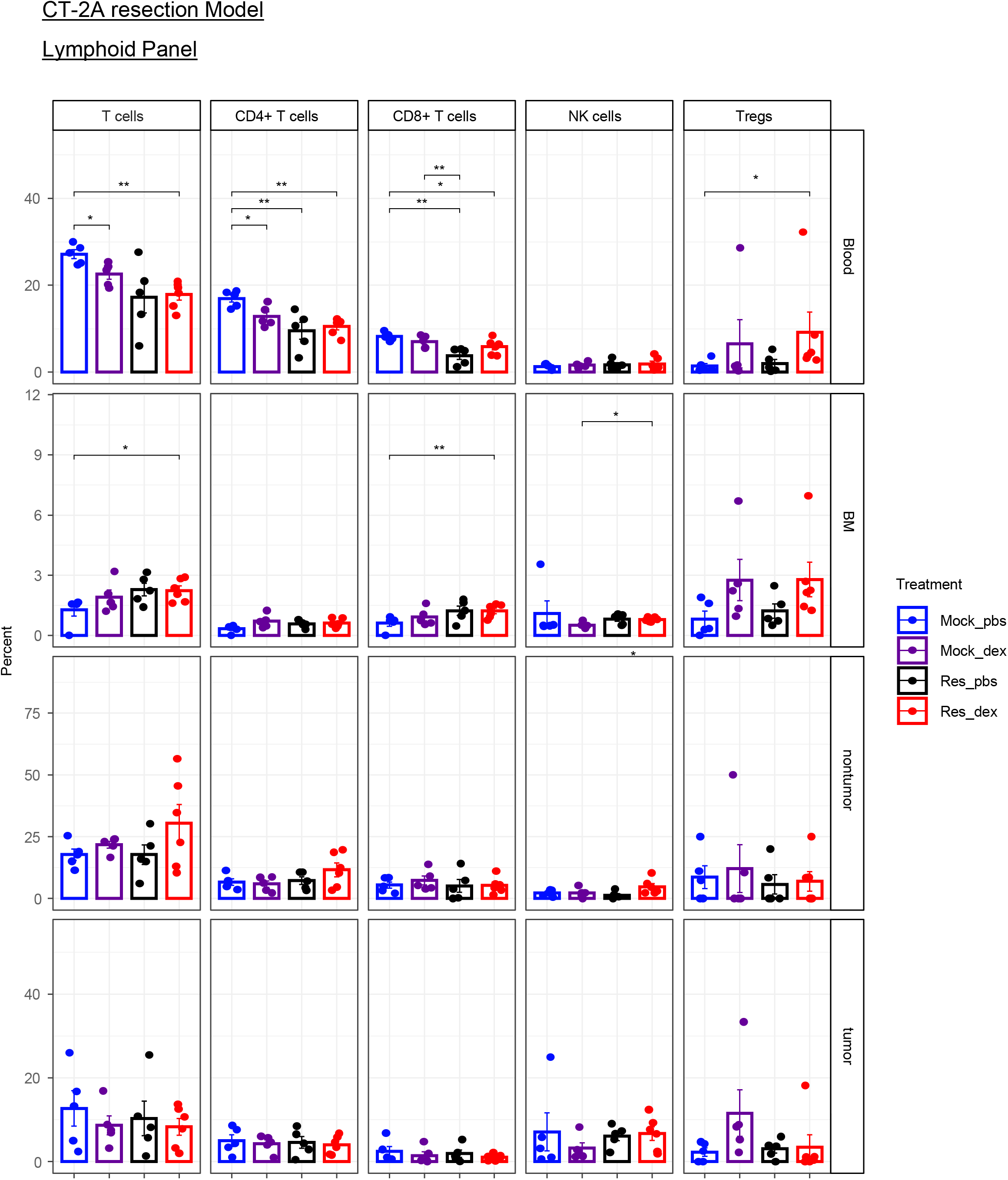
CT-2A-bearing mice as described in Figure 1 and Figure 2, including the groups mock PBS, mock dexamethasone, resection PBS, and resection dexamethasone, were evaluated via flow cytometry for T cells, CD4+ T cells, CD8+ T cells, NK cells, and T-regulatory cells in the blood, bone marrow, non-tumor cortex, spleen, and tumor. Groups were compared by t-tests; *p<0.05, **p<0.01, ***p<0.001.

**Supplemental Figure 8.**
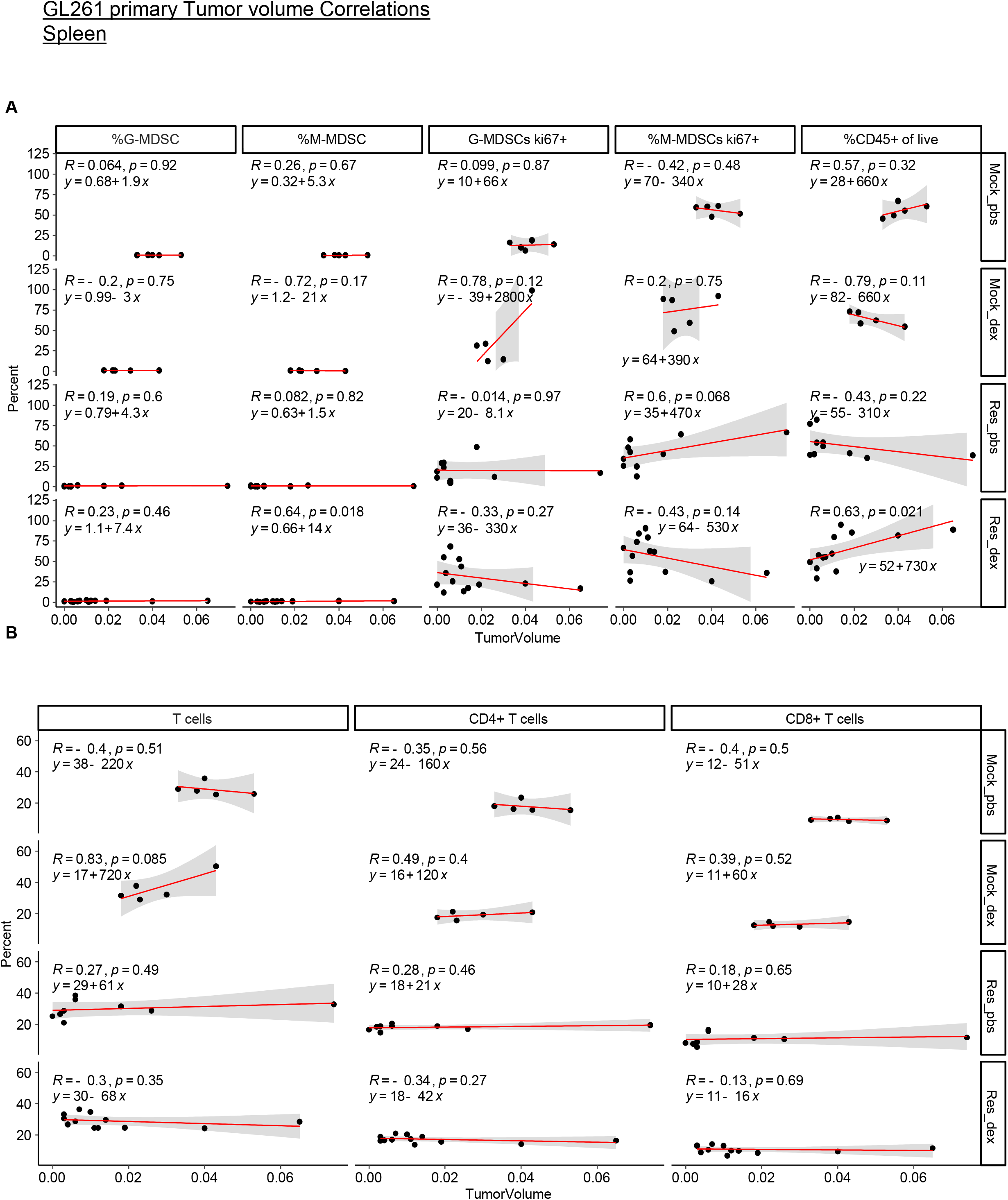
The spleen myeloid populations and total CD45+ cells of live cells (**A**) and splenic T cell populations (**B**) are shown for n=10 mock PBS, n=9 mock dexamethasone, n=14 resection PBS, and n=13 resection dexamethasone mice. This corresponds to data included in Figure 3 showing the tumor volume correlation with splenic myeloid populations via flow cytometry. Correlation coefficients (R), p values and fitted line parameters are shown.

**Supplemental Figure 9.**
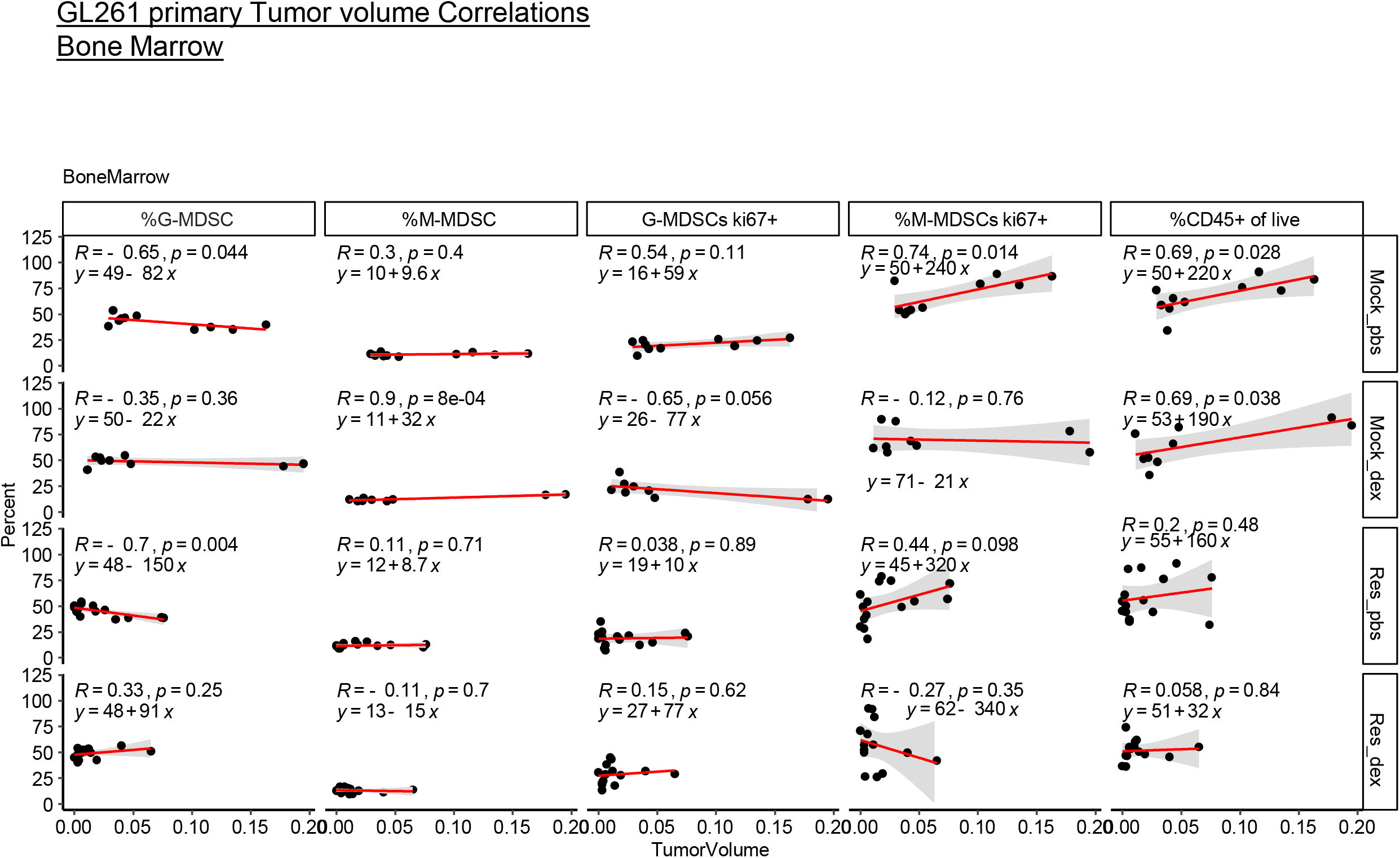
The bone marrow myeloid populations and total CD45+ cells of live cells are shown for n=10 mock PBS, n=9 mock dexamethasone, n=14 resection PBS, n=13 resection dexamethasone mice. This corresponds to data included in Figure 3 data showing the tumor volume correlation with splenic myeloid populations via flow cytometry. Correlation coefficients (R), p values and fitted line parameters are shown.

**Supplemental Figure 10.**
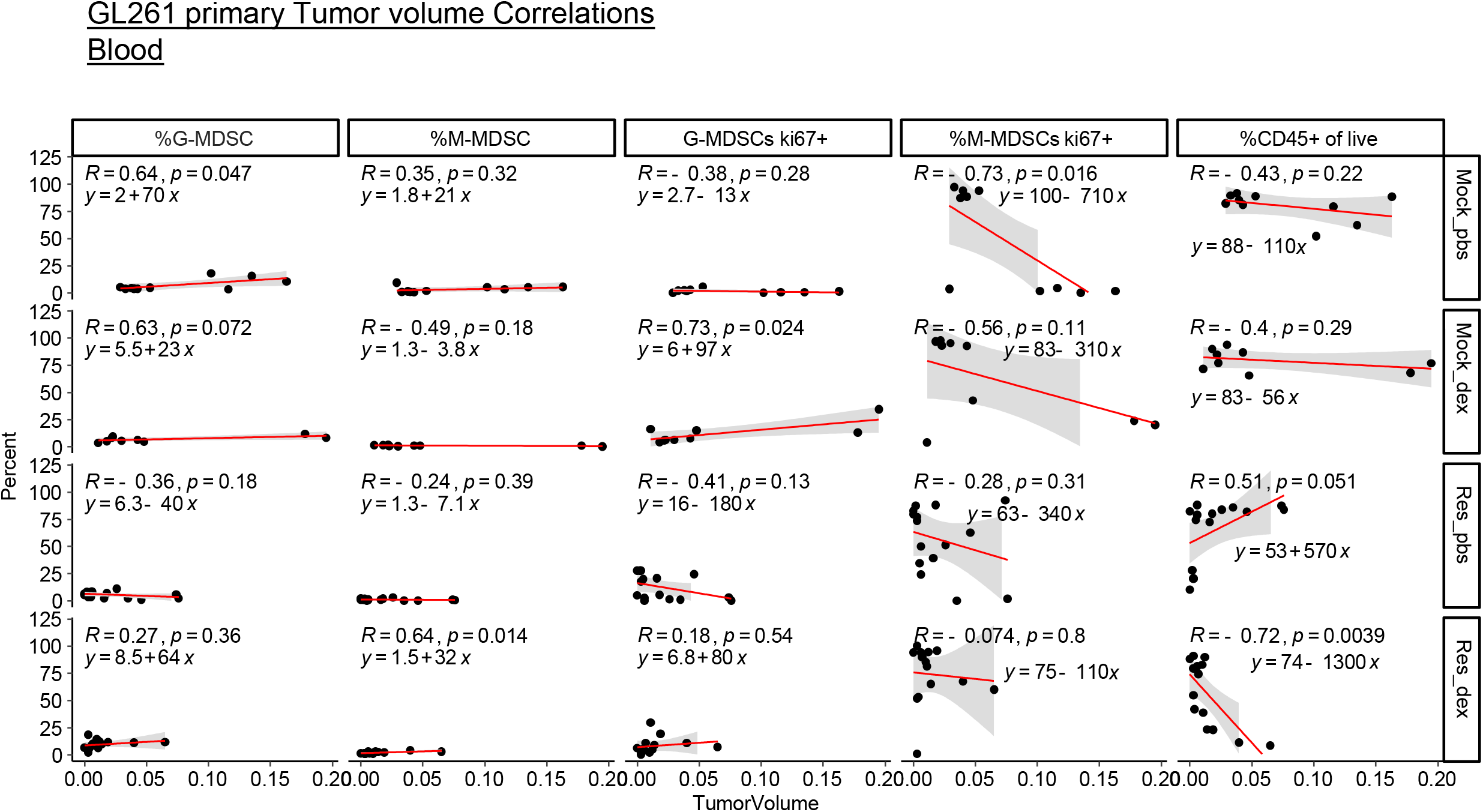
The blood-derived myeloid populations and total CD45+ cells of live cells are shown for n=10 mock PBS, n=9 mock dexamethasone, n=14 resection PBS, n=13 resection dexamethasone mice. This corresponds to data included in Figure 3 data showing the tumor volume correlation with splenic myeloid populations via flow cytometry. Correlation coefficients (R), p values and fitted line parameters are shown.

**Supplemental Figure 11.**
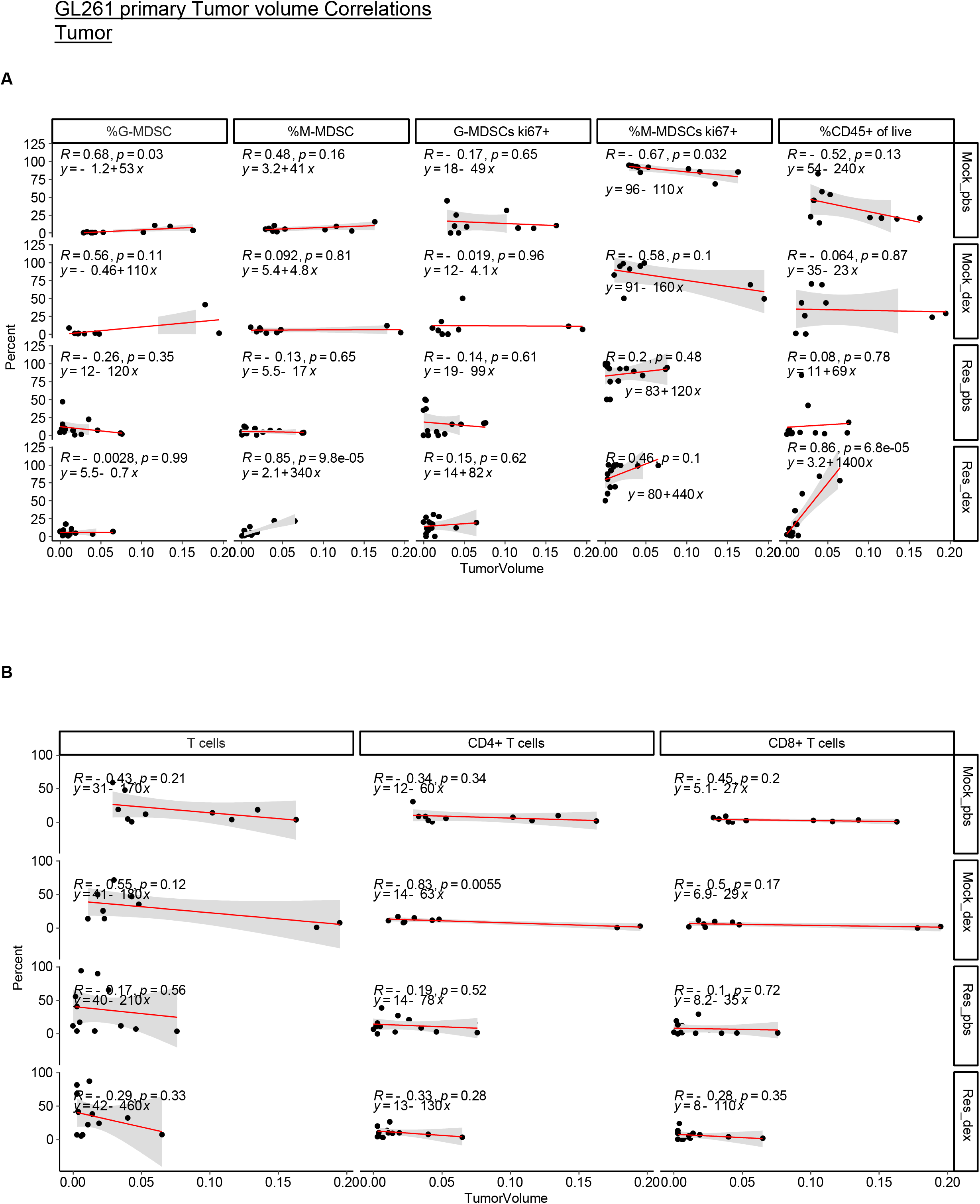
The tumor-derived myeloid populations and total CD45+ cells of live cells (**A**) along with T cell populations (**B**) are shown for n=10 mock PBS, n=9 mock dexamethasone, n=14 resection PBS, n=13 resection dexamethasone mice. This corresponds to data included in Figure 3 data showing the tumor volume correlation with splenic myeloid populations via flow cytometry. Correlation coefficients (R), p values and fitted line parameters are shown.

**Supplemental Figure 12.**
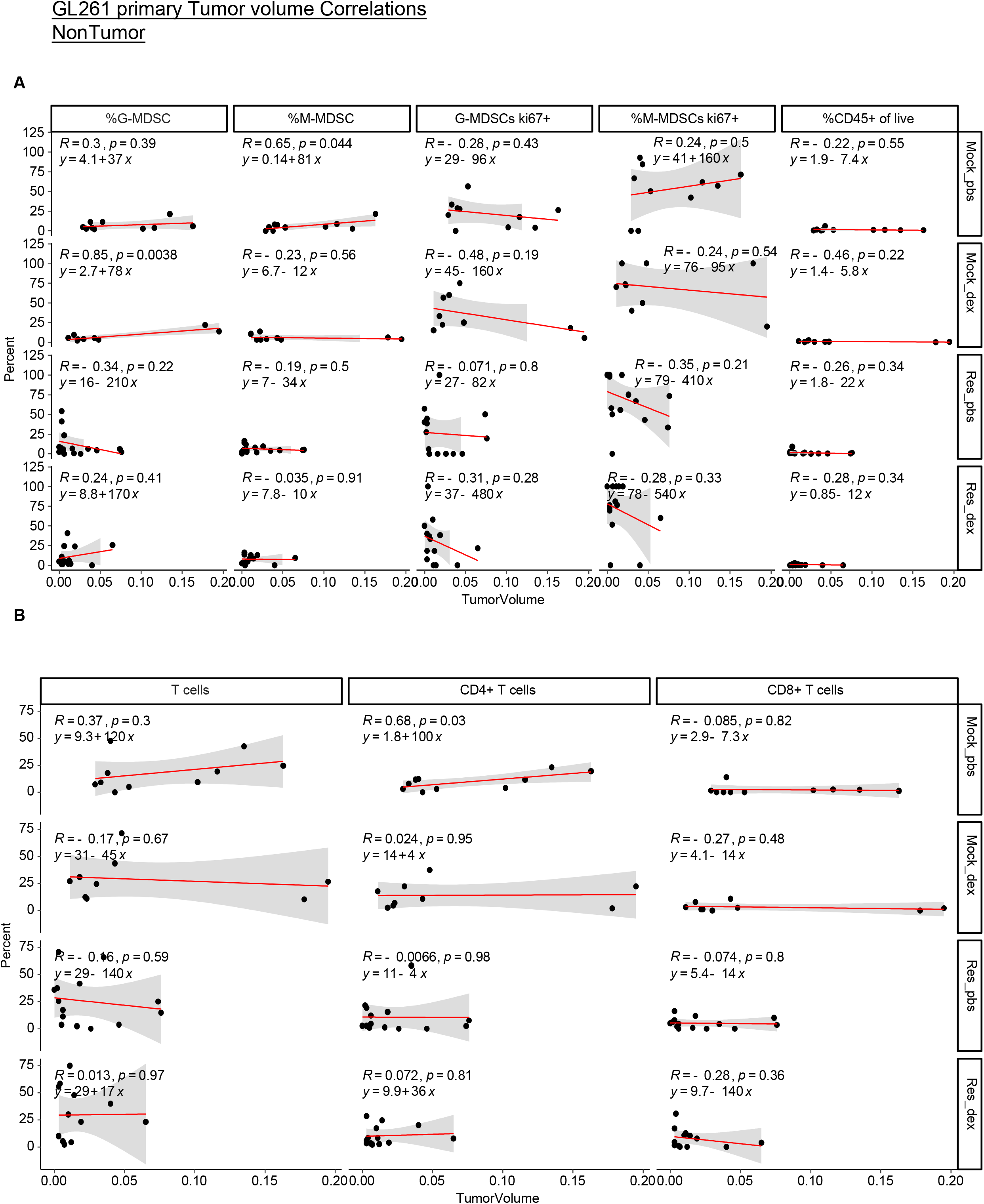
The non-tumor cortex-derived myeloid populations and total CD45+ cells of live cells (**A**) along with T cell populations (**B**) are shown for n=10 mock PBS, n=9 mock dexamethasone, n=14 resection PBS, n=13 resection dexamethasone mice. This corresponds to data included in Figure 3 showing the tumor volume correlation with splenic myeloid populations via flow cytometry. Correlation coefficients (R), p values and fitted line parameters are shown.

**Supplemental Figure 13.**
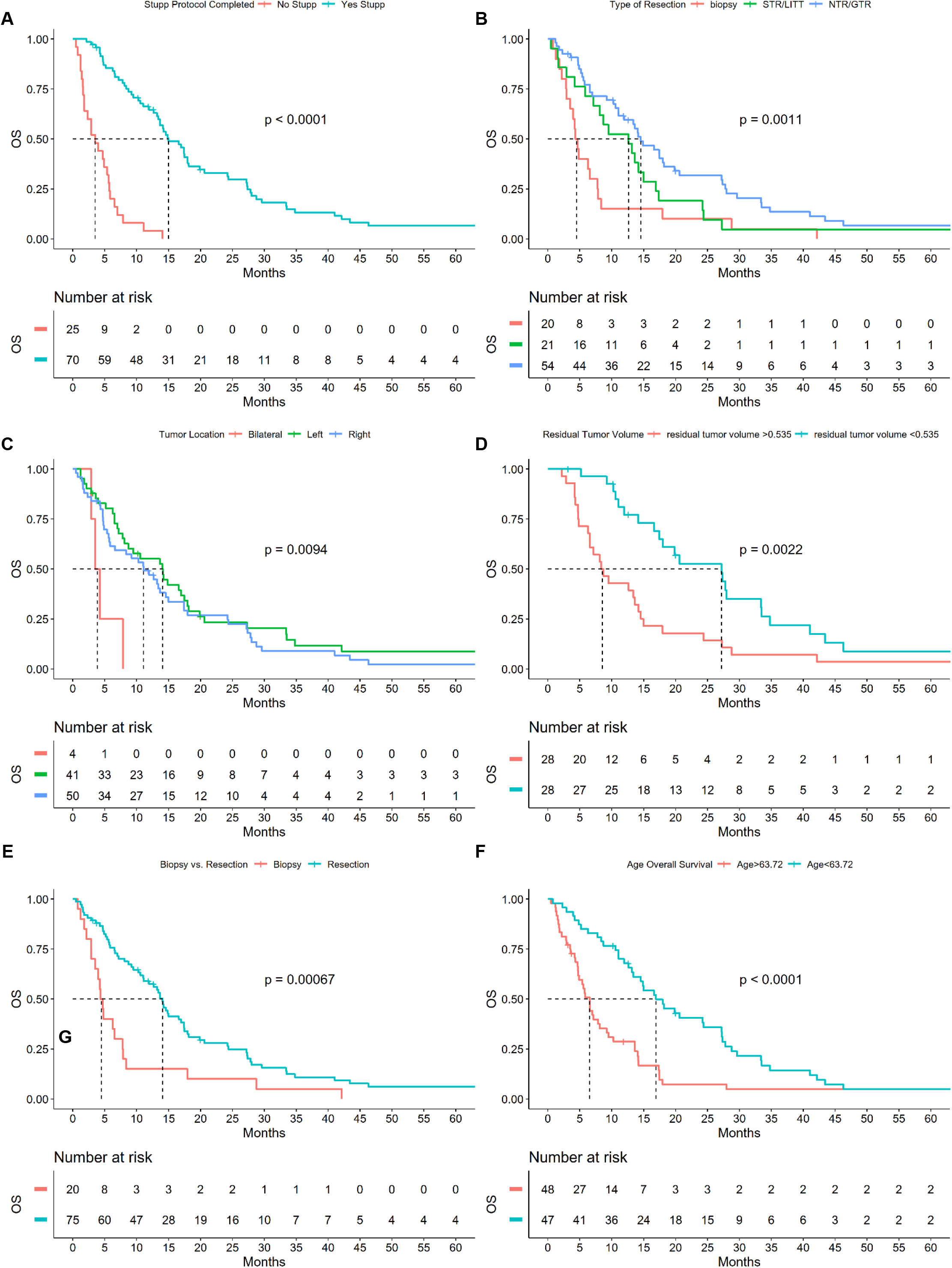
Univariate Kaplan Meier analysis of overall survival GBM cohort n=95 patients. Kaplan Meier comparing those who completed Stupp Protocol vs those who did not (**A**). Kaplan Meier comparing those who had a Biopsy vs subtotal resection/LITT therapy, vs near total or gross total resection (**B**). Kaplan Meier comparing the tumor location L=Left hemisphere, R=Right hemisphere, B=Bilateral (**C**). Kaplan Meier comparing the residual tumor volume post resection, divided by median 0.535 (**D**). Kaplan Meier comparing those who had Biopsy vs resection of any type (**E**). Kaplan Meier comparing age split by median 63.72 years (**F**). All P values represent log rank comparison and dotted lines represent the median survival times for each curve.

**Supplemental Figure 14.**
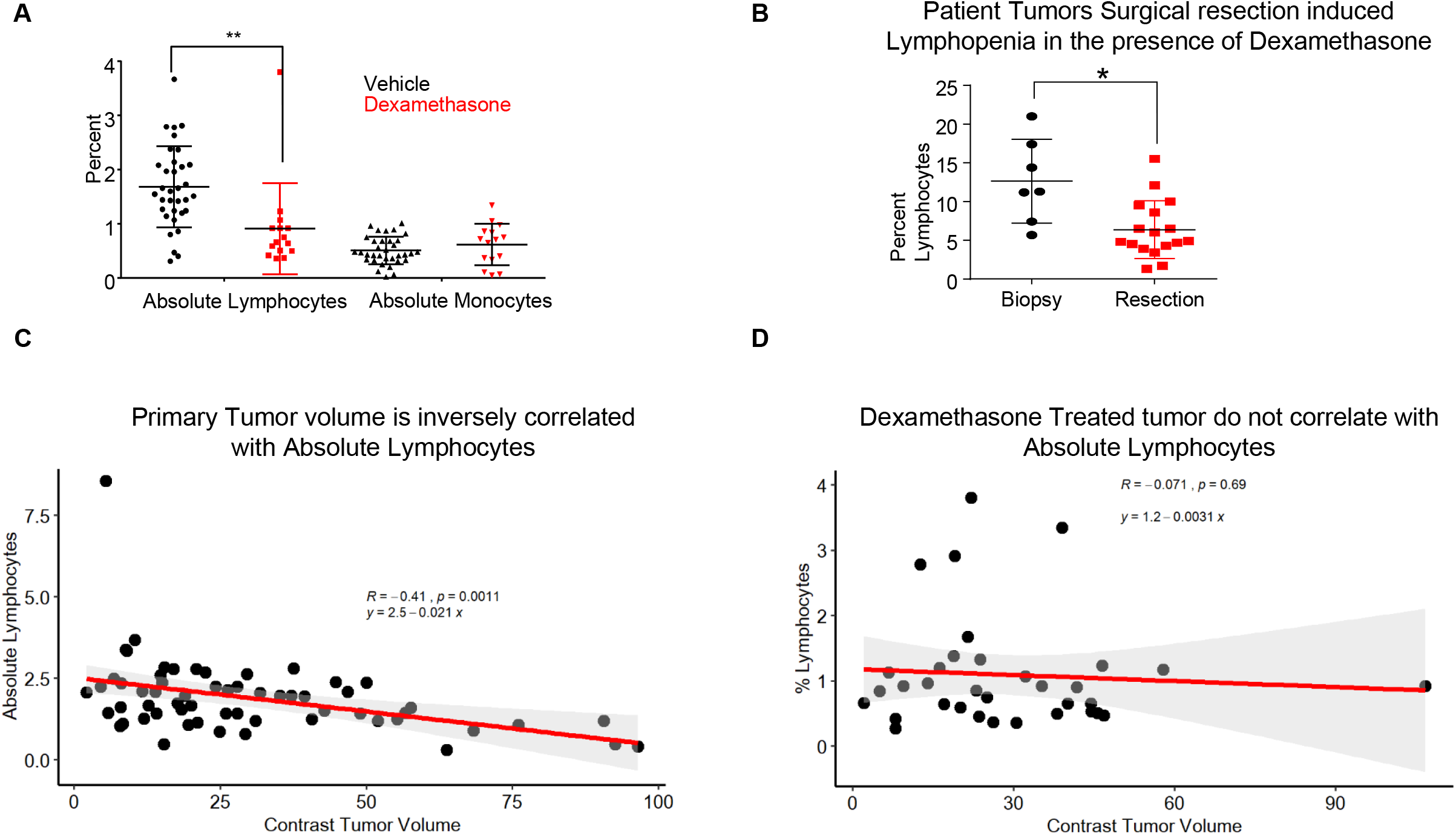
Corresponding to Figure 4, absolute lymphocyte count vs tumor volume was graphed for patients prior to surgery or other treatment (**A**). Similarly, lymphocyte levels post-intervention were graphed for dexamethasone-treated patients, prior to surgery or biopsy (**B**). Corresponding to Figure 4, the absolute lymphocytes were graphed against tumor volume in steroid-naïve and steroid-treated patients (**C, D**). Correlation coefficients (R), p values and fitted line parameters are shown. Groups were compared by t-tests; *p<0.05, **p<0.01, ***p<0.001.

**Supplemental Figure 15.**
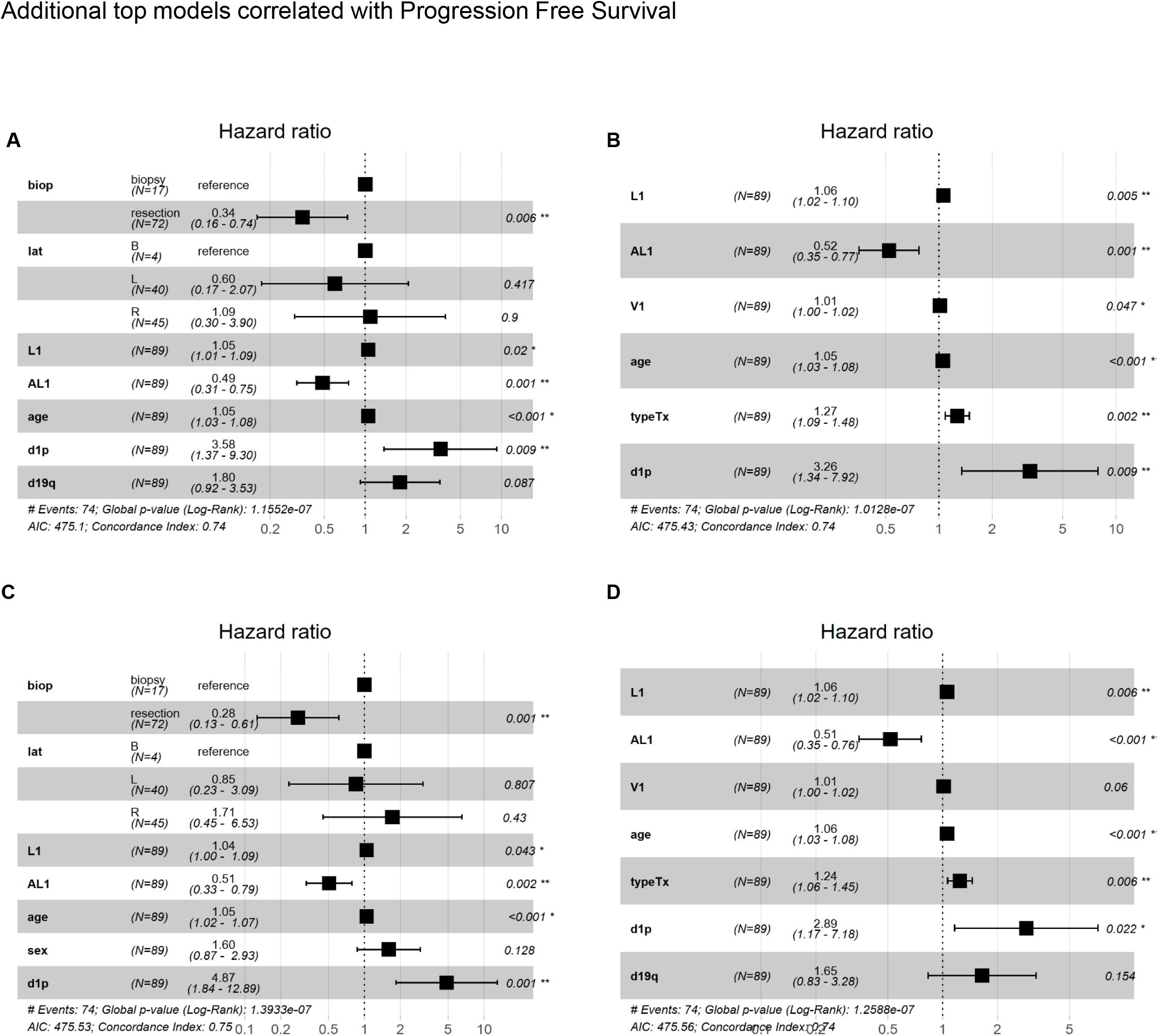
Cox proportional hazards models of progression free survival automatically selected based on AIC using R package glmulti. In this analysis the top model is in **Fig 4D** and the next 4 are in (**A-D**) here. Biop=Biopsy vs resection, lat=Tumor Laterality, L1= % lymphocytes pre surgery, AL1= Absolute lymphocyte count pre surgery, d1p=chromosome 1p status, d19q=19q status, V1= tumor volume at diagnosis, typeTx= type of tumor resection.

**Supplemental Figure 16.**
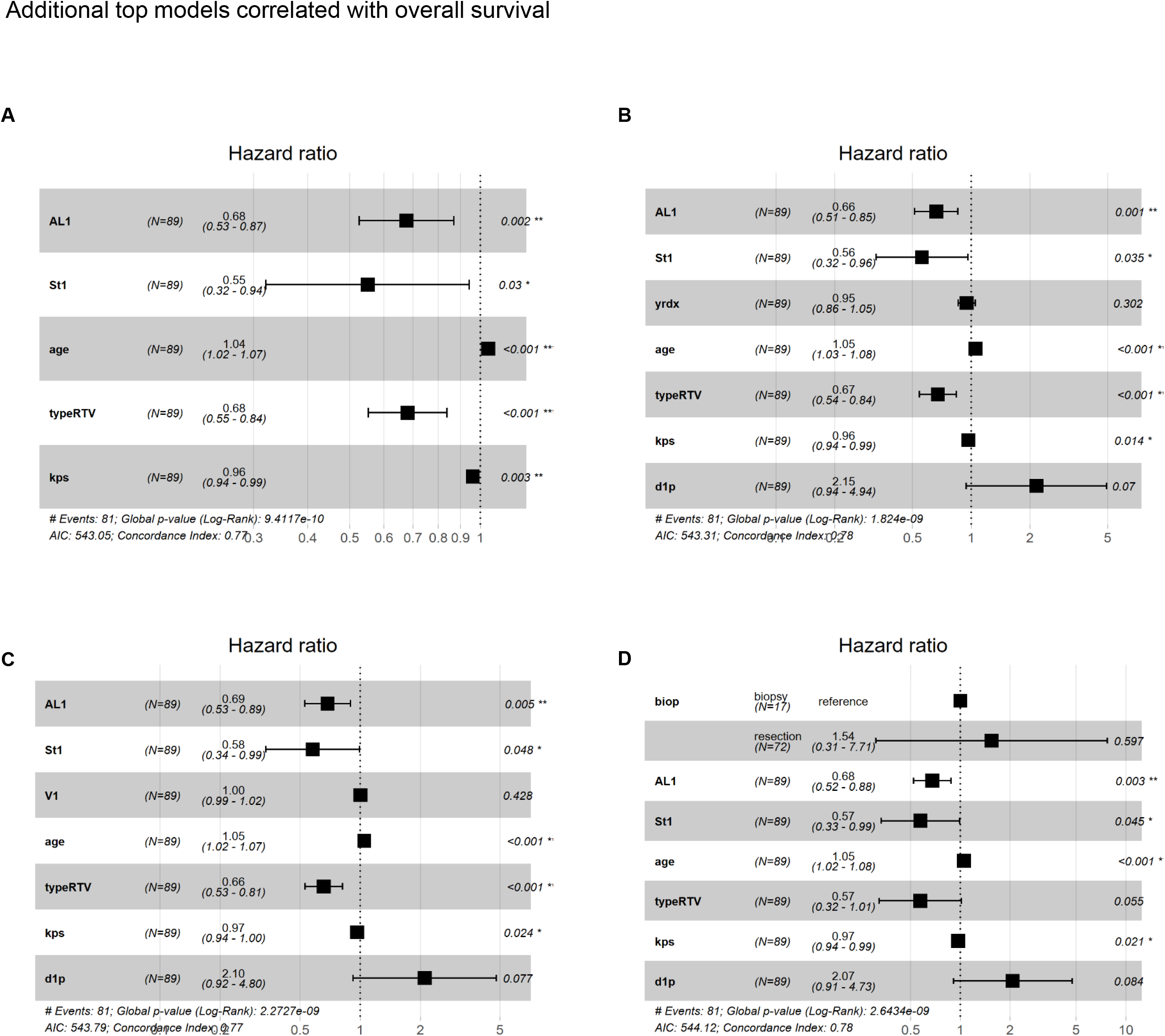
Cox proportional hazards models of overall survival automatically selected based on AIC using R package glmulti. In this analysis the top model is represented in **Fig 4E**, the additional 4 models identified using this method are represented in (**A-D**). Biop=Biopsy vs resection, St1= Steroids pre op at time of lymphocyte measurement, typeRTV= Type of tumor resection, L1= % lymphocytes pre surgery, AL1= Absolute lymphocyte count pre surgery, d1p=chromosome.

**Supplemental Figure 17.**
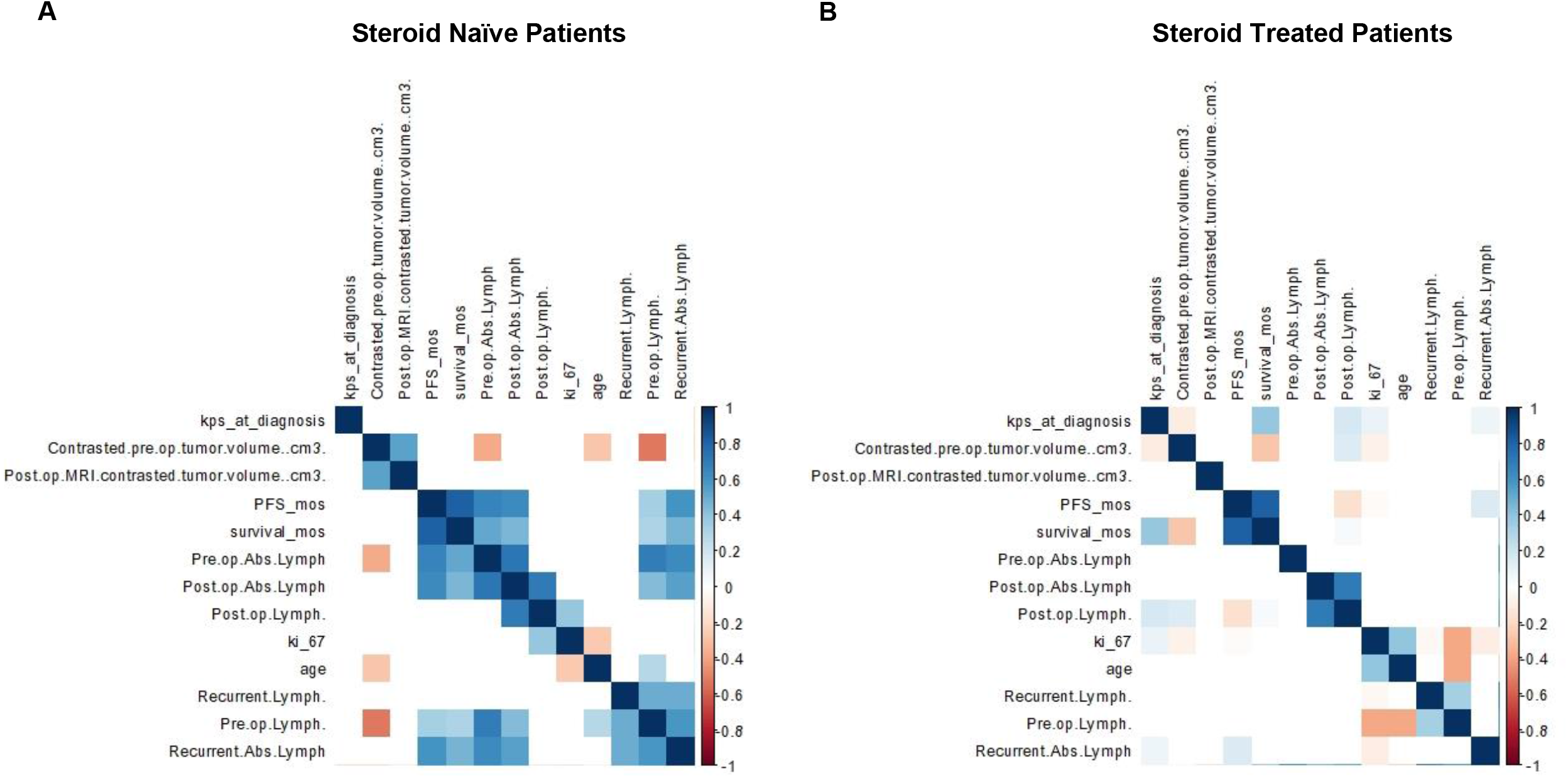
Inter-variable correlation showing variable co-dependence when steroid-treated patients vs non-steroid treated patients n=61 and n=34 respectively (**A-B**). Correlation plots only show correlations with p<0.05 with the correlation coefficient colored by the scale −1 to 1 to the right of plot.

