## Supplementary figures and images for "Preclinical modeling of surgery and steroid therapy for glioblastoma reveals changes in immunophenotype that are associated with tumor growth and outcome"

### Supplemental Fig. 1

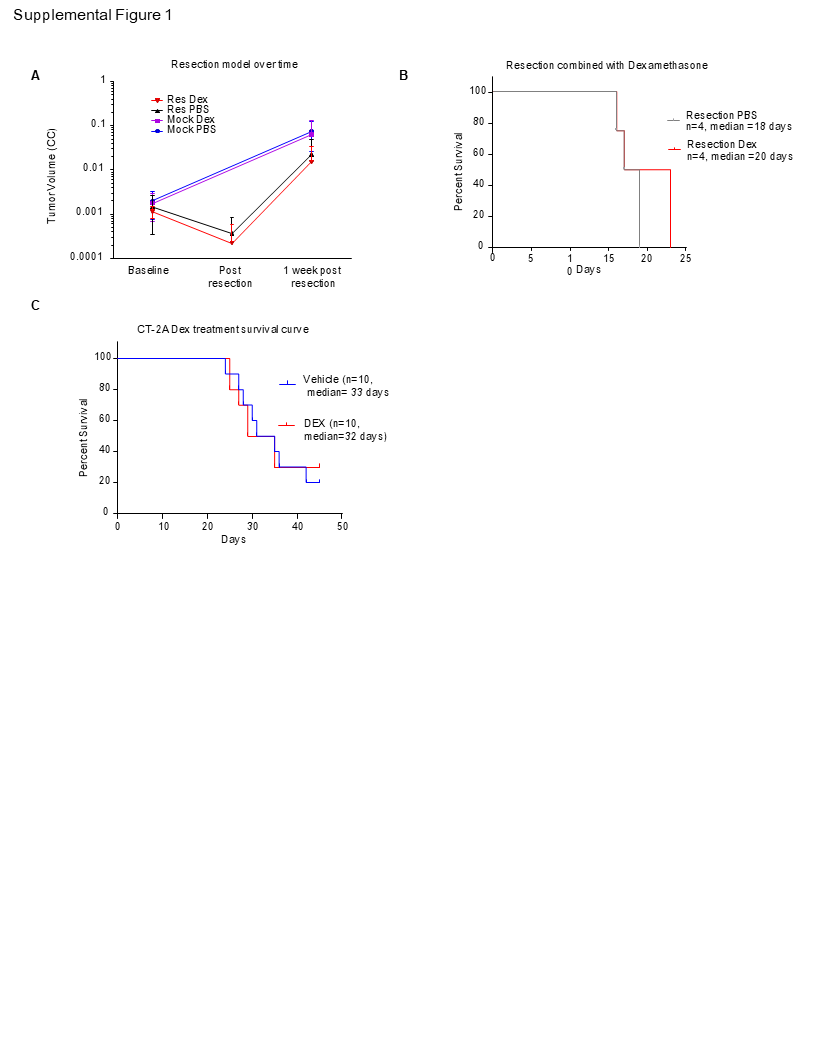

### Supplemental Fig. 2

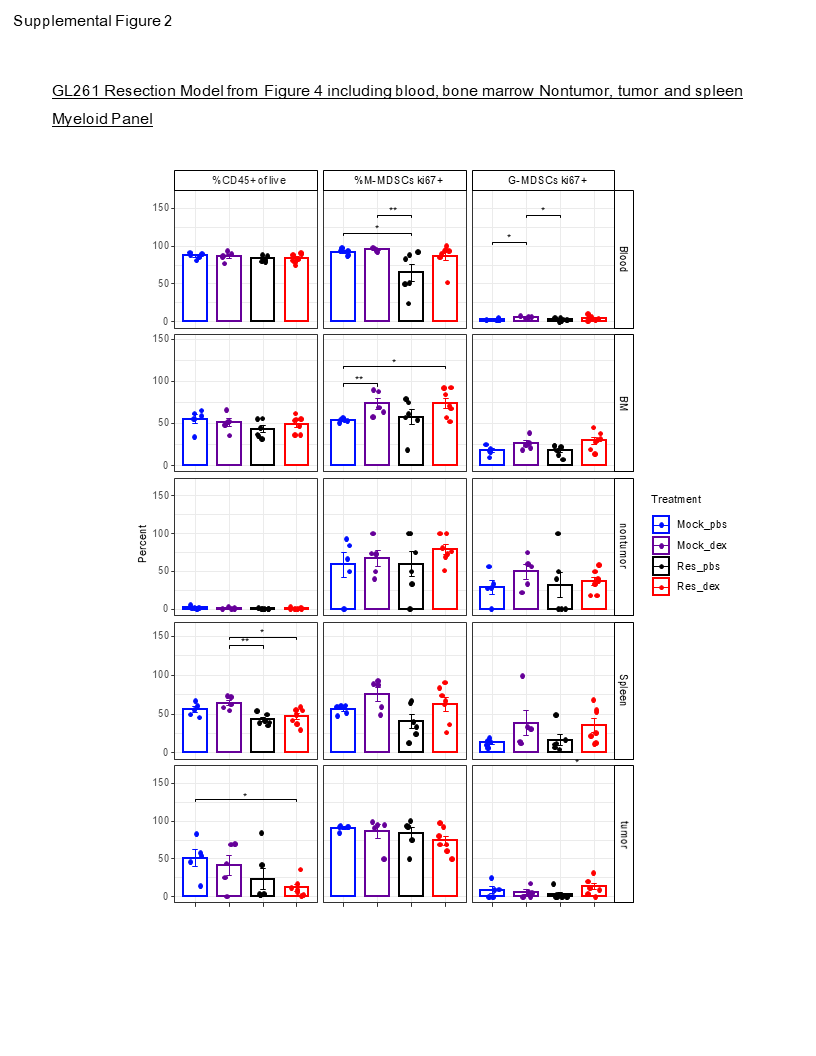

### Supplemental Fig. 3

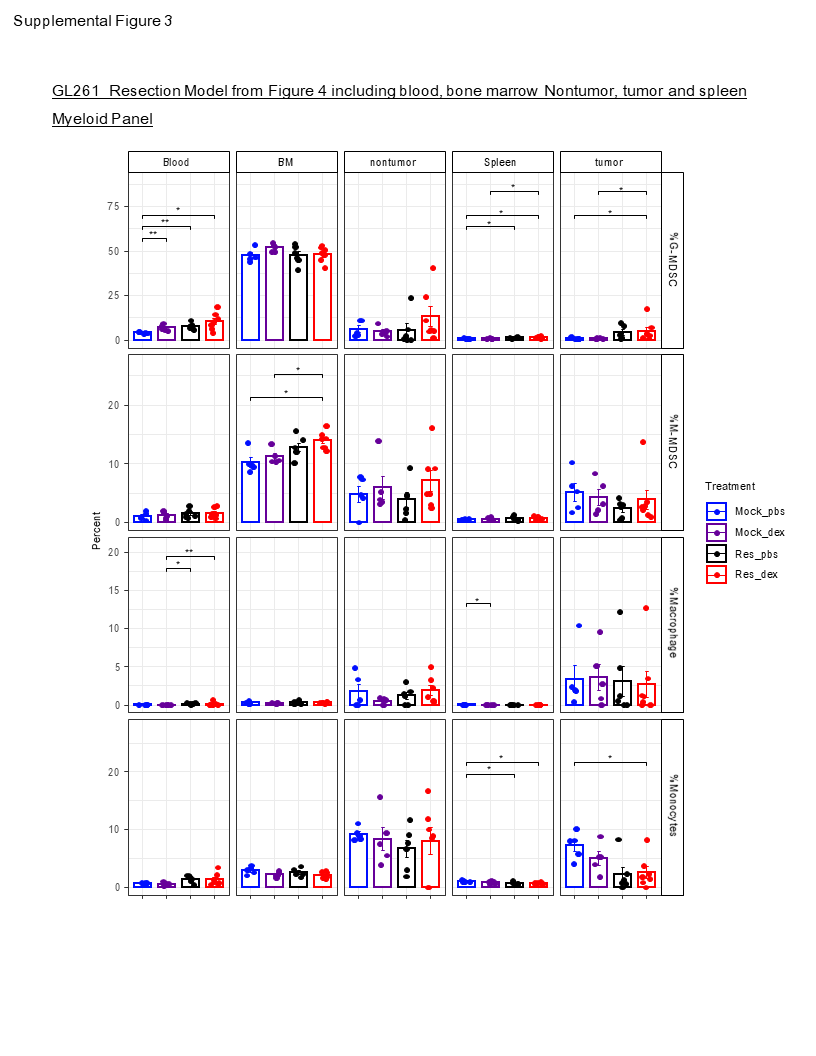

### Supplemental Fig. 4

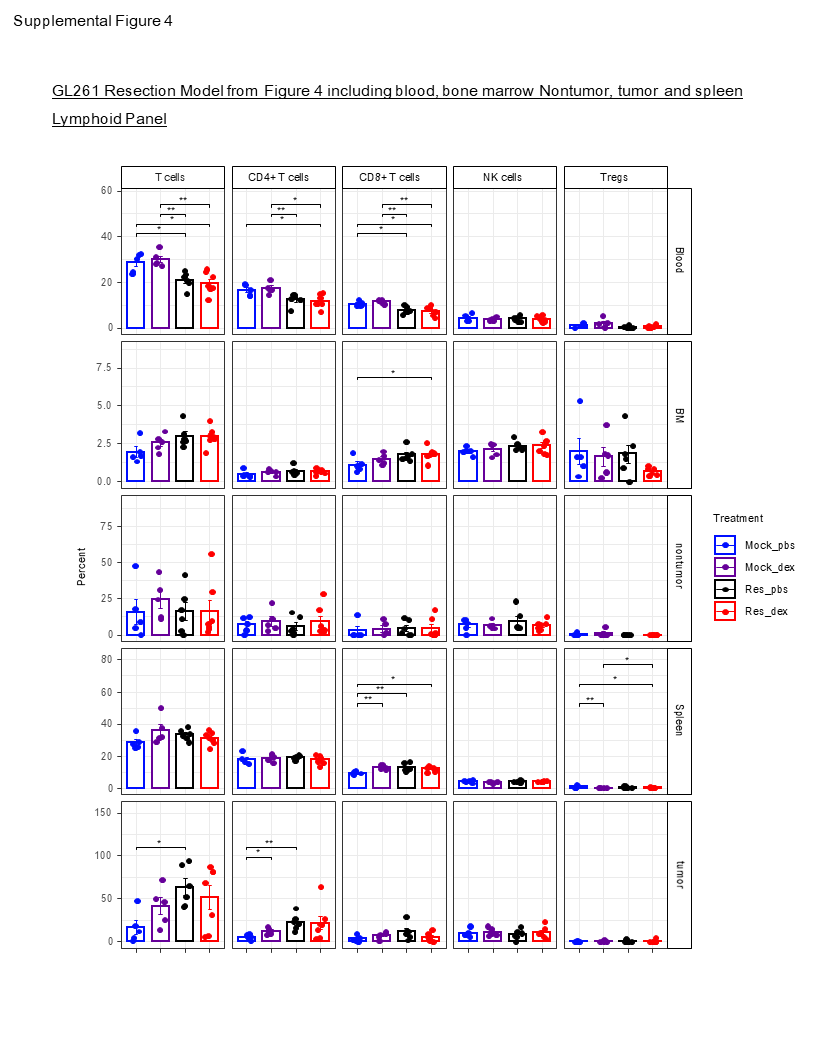

### Supplemental Fig. 5

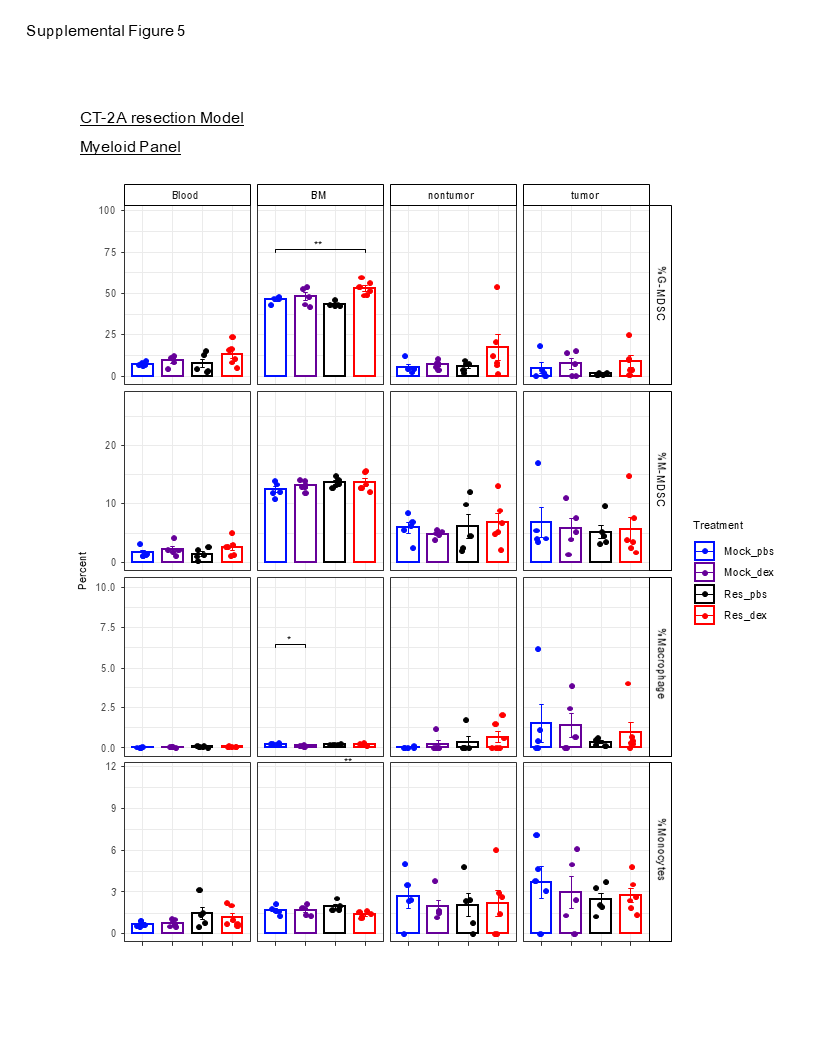

### Supplemental Fig. 6

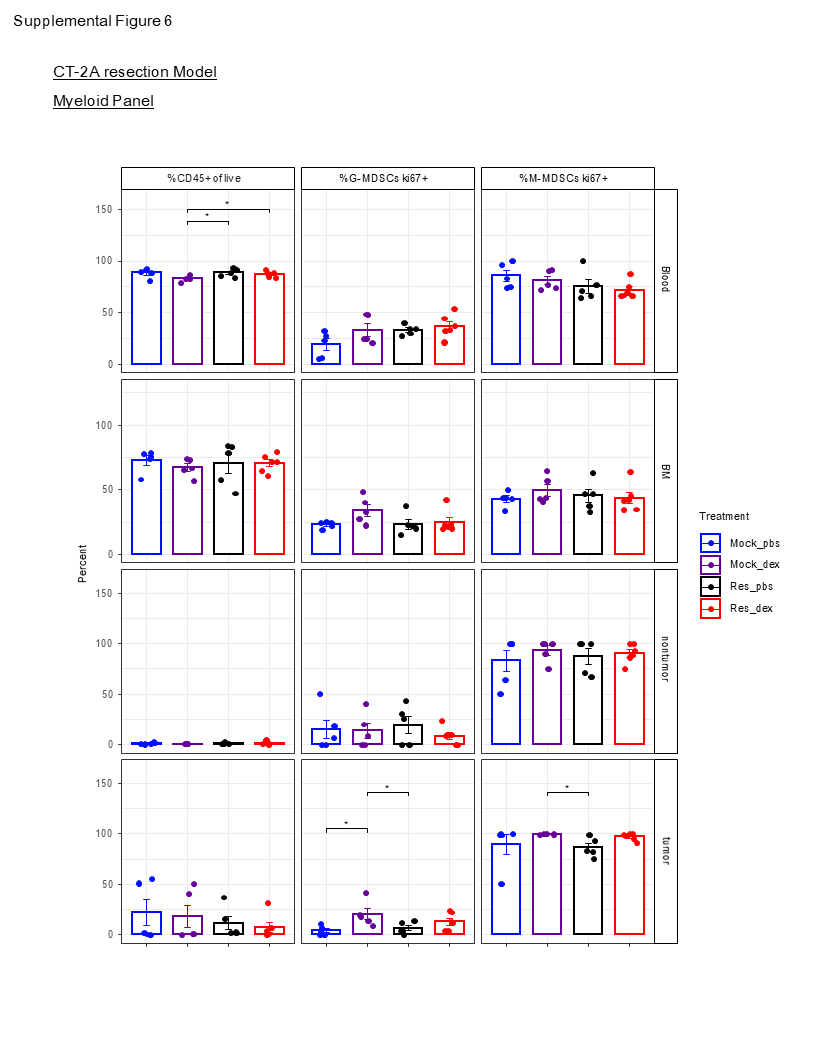

### Supplemental Fig. 7

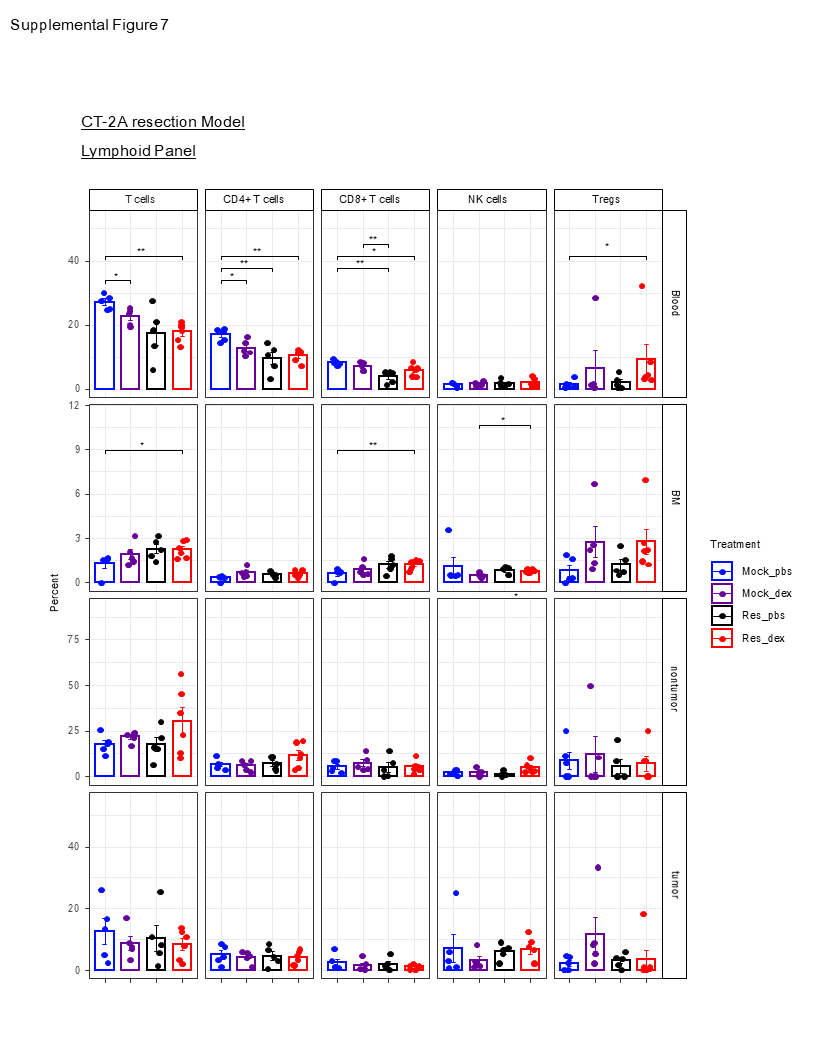

### Supplemental Fig. 8

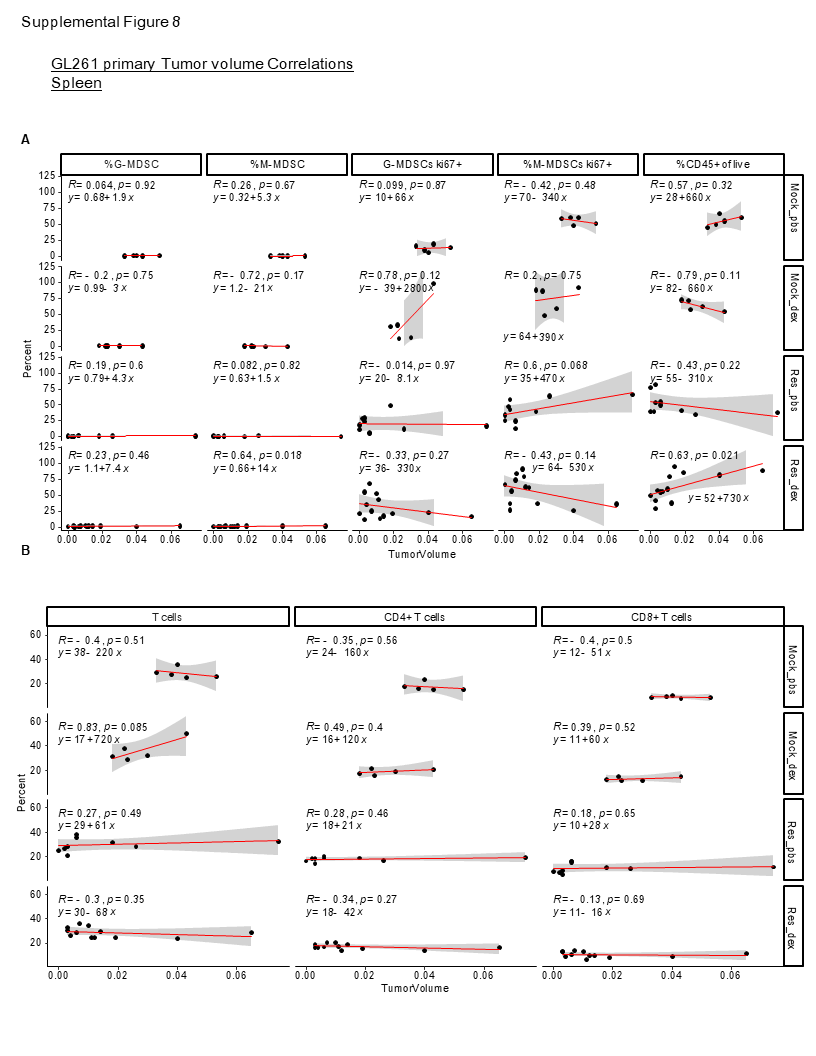

### Supplemental Fig. 9

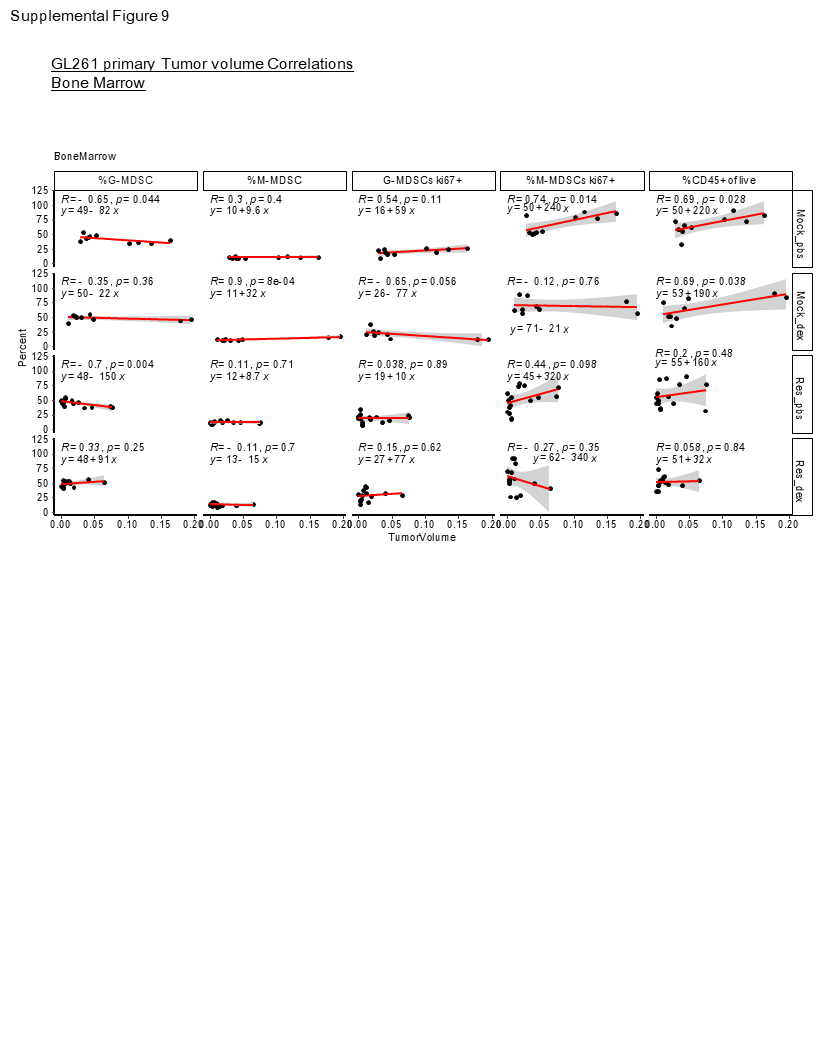

### Supplemental Fig. 10

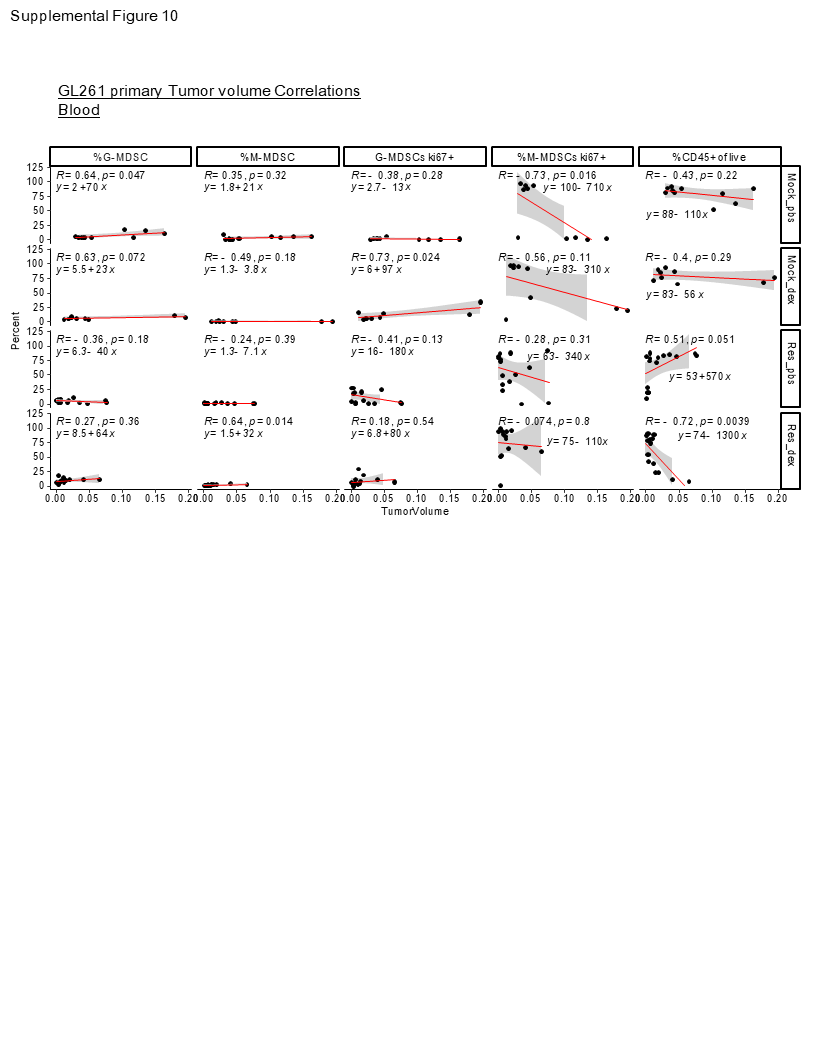

### Supplemental Fig. 11

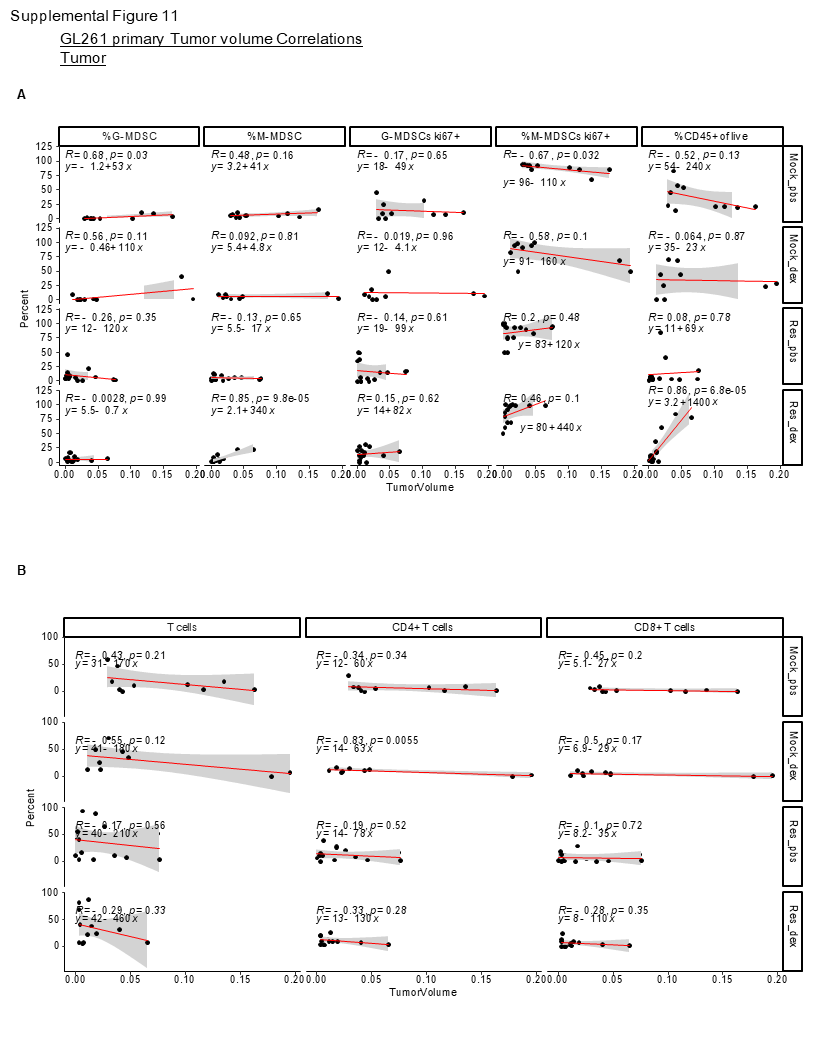

### Supplemental Fig. 12

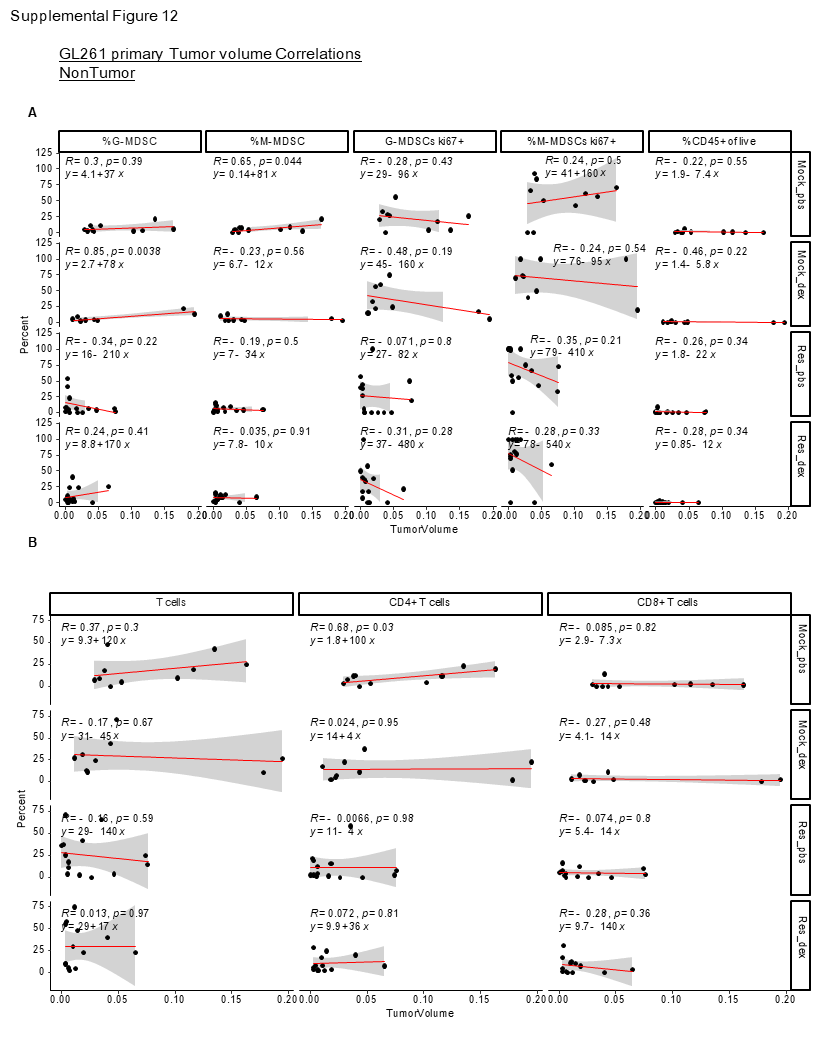

### Supplemental Fig. 13

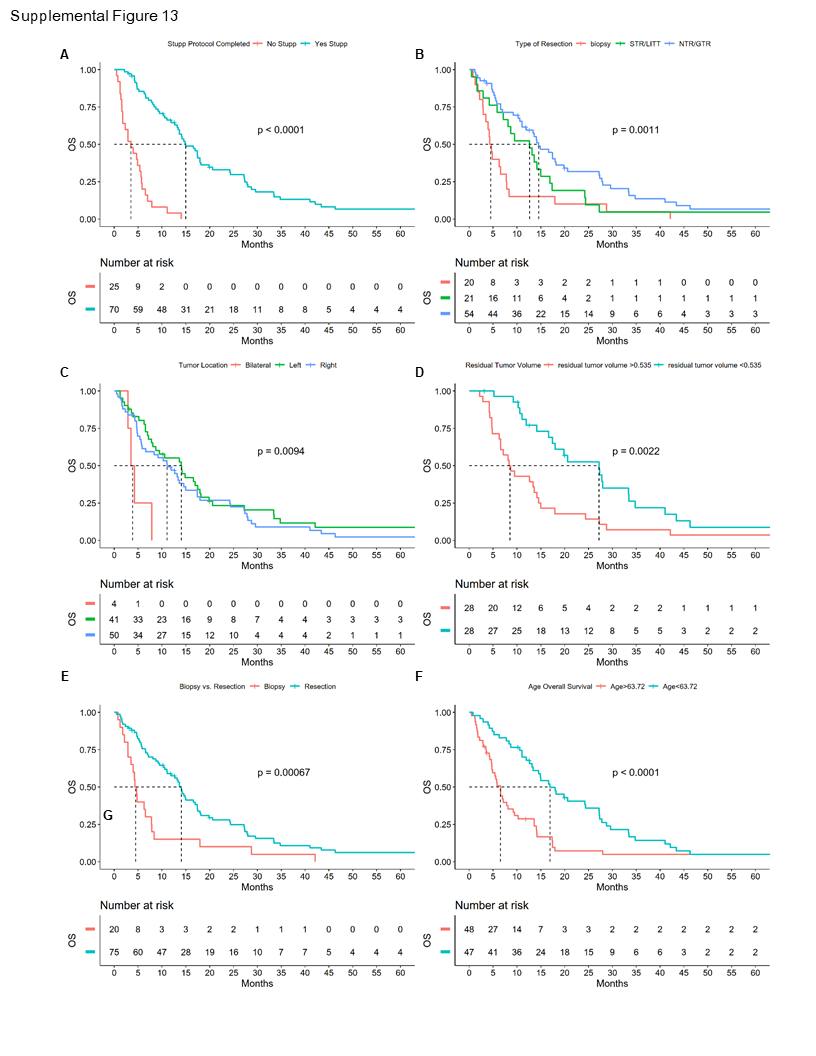

### Supplemental Fig. 14

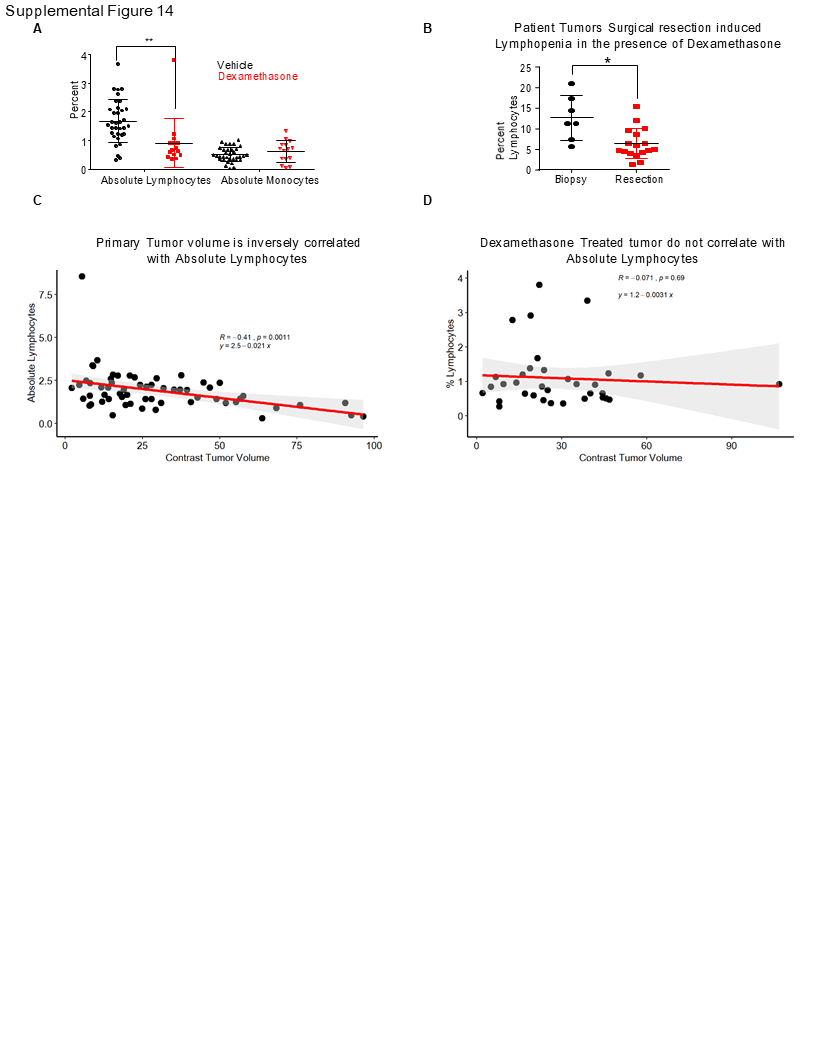

### Supplemental Fig. 15

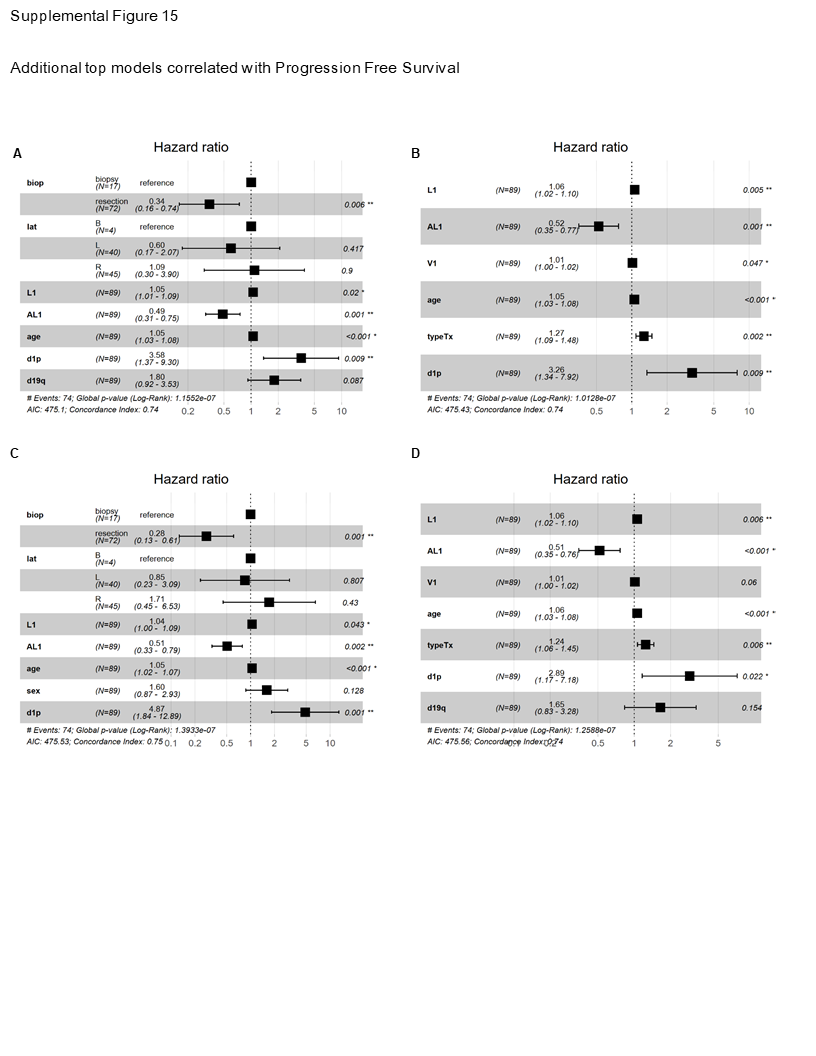

### Supplemental Fig. 16

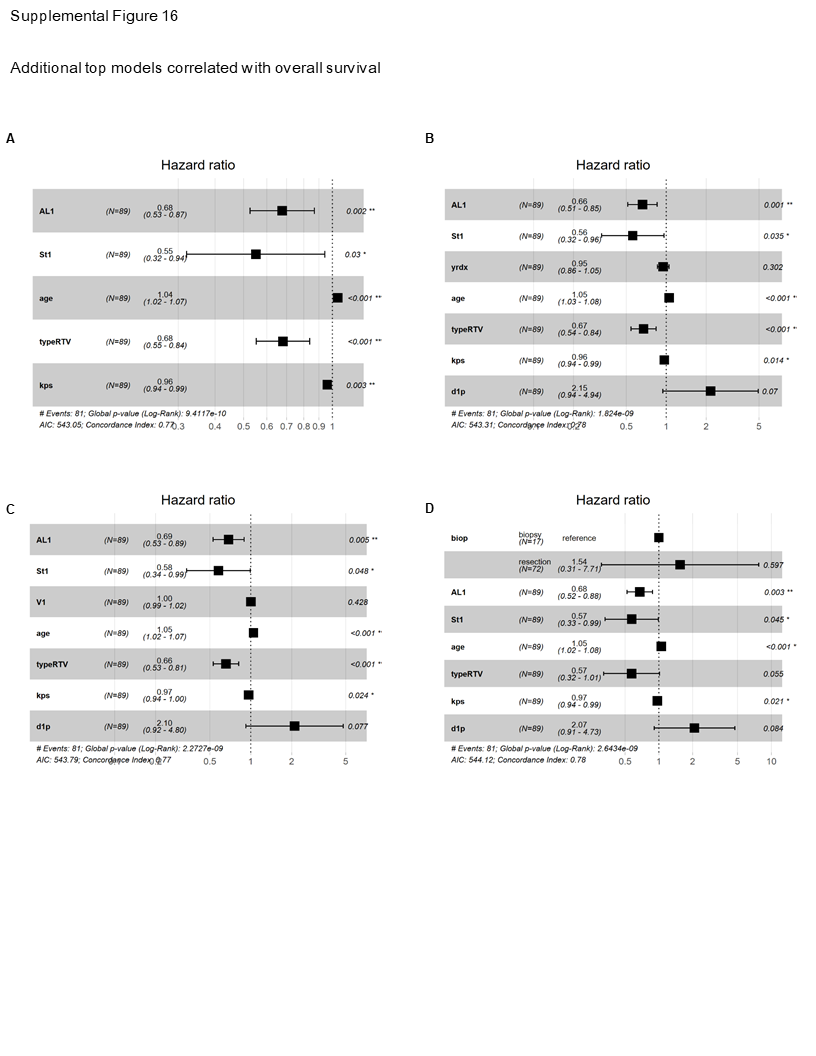

### Supplemental Fig. 17

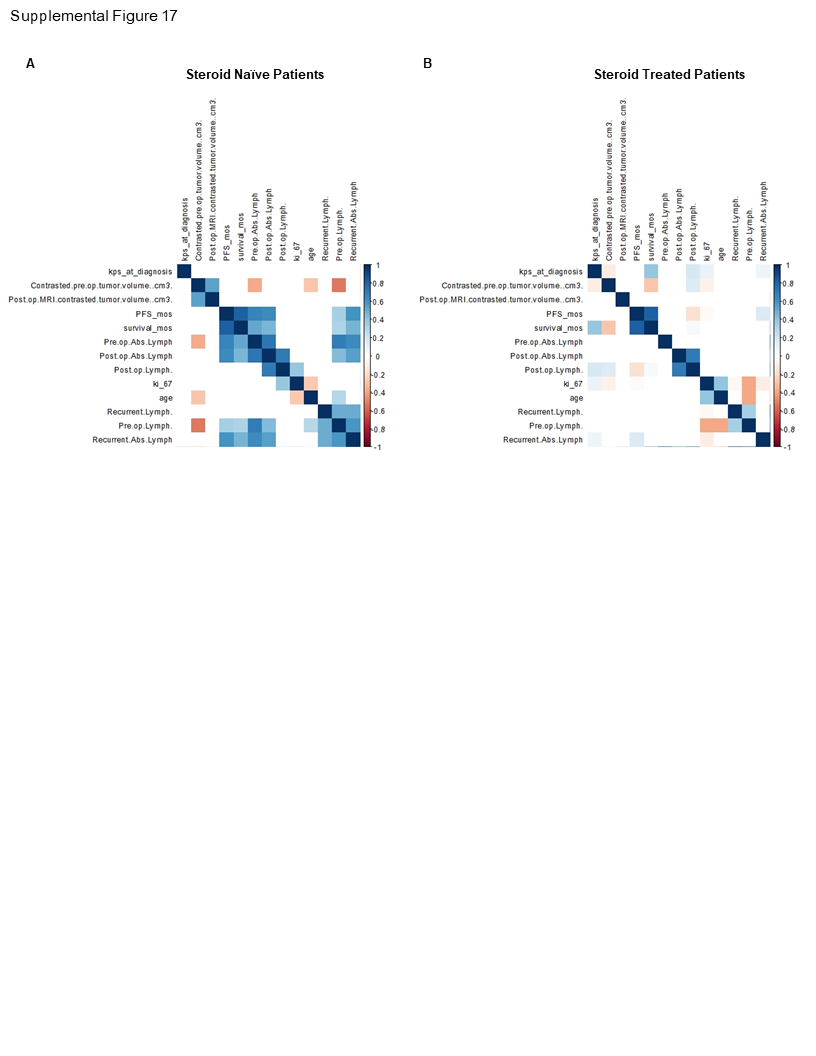
